# Benchmarking computational tools for locus-specific analysis of transposable elements in single-cell RNA-seq datasets

**DOI:** 10.64898/2026.02.26.708244

**Authors:** Veronica Finazzi, Catalina A. Vallejos, Antonio Scialdone

**Affiliations:** Institute of Epigenetics and Stem Cells, Helmholtz Zentrum München, Munich, Germany; Institute of Computational Biology, Helmholtz Zentrum München, Neuherberg, Germany; Institute of Functional Epigenetics, Helmholtz Zentrum München, Neuherberg, Germany; Institute of Genetics and Cancer, University of Edinburgh, Edinburgh, UK; Health Data Research UK, London, UK

**Keywords:** transposable elements, single-cell RNA-seq, locus-specific quantification, benchmarking, computational

## Abstract

**Background:** Transposable elements (TEs) are increasingly recognized as regulators of gene expression and cellular identity in development and disease. Single-cell RNA-sequencing (scRNA-seq) enables the analysis of their transcription at cellular resolution, but the repetitive nature of TEs and their frequent overlap with genes create substantial mapping ambiguity. Although several tools quantify TE expression, few support locus-specific analysis, and their performance in single-cell data has not been systematically evaluated.

**Results:** We present a comprehensive benchmarking framework for locus-level TE quantification in short-read scRNA-seq, combining real datasets with simulations that provide read-level ground truth. TE-derived reads constitute a considerable fraction of the transcriptome and capture meaningful biological structure. Our simulations reveal that older, sequence-diverged insertions can be quantified with relatively high accuracy, whereas young TEs remain intrinsically difficult to resolve due to unreliable assignment of multi-mapping reads. We observe pronounced family-specific biases and identify gene-TE disambiguation as a major unresolved challenge. Among evaluated methods, SoloTE (unique-mapper mode) and Stellarscope (with an expectation-maximization-based reallocation of multi-mappers) showed comparable performance, while including multi-mappers generally increased false positives without substantially improving locus-level accuracy.

**Conclusions:** Our benchmark delineates the fundamental limits imposed by short-read scRNA-seq on locus-specific TE quantification, providing practical guidance for prospective users. Suggested best practices include focusing locus-level analyses on older insertions, applying unique-mapper strategies to improve precision, aggregating counts at the subfamily level for young TEs, and explicitly checking for gene-TE overlaps. Our workflow is fully reproducible and extensible, providing a foundation for evaluating emerging methods aimed at resolving TE transcription at single-locus resolution.

## 1 Introduction

Transposable elements (TEs) are increasingly recognized as major contributors to gene regulation and cellular identity across diverse biological contexts, including embryonic development, neurodegerative diseases and cancer [1, 2, 3, 4]. Although most TE insertions have lost mobilization capacity throughout evolution, the sequences they leave behind can act as regulatory elements or give rise to regulatory RNAs with important cellular functions [5, 6, 7, 8]. TE activity is highly celltype and stage specific: while most loci remain epigenetically repressed, a small subset is activated in a tightly regulated manner [9, 10, 11]. Yet, the mechanisms controlling which loci are active and how specific insertions influence neighboring genes or cellular programs remain unclear [12, 9, 10]. Single-cell omics technologies offer a unique opportunity to study TE activity with the resolution required to capture its cellular heterogeneity and dynamic changes, which are obscured in bulk measurements. For example, waves of TE activation during mammalian preimplantation development are essential for embryonic genome activation and the acquisition of totipotent identity [2]. Nevertheless, recent single-cell atlases of early embryos continue to exclude TEs from their analyses [13, 14], reflecting a broader trend in which TEs remain systematically overlooked in standard scRNA-seq workflows.

A major reason for this omission is the technical difficulty of quantifying TE expression [15]. TEs are repetitive and interspersed throughout the genome. Sequencing reads derived from them frequently map to multiple loci, complicating assignment. Overlaps between TEs and genes can generate chimeric or ambiguous transcripts, making it difficult to distinguish autonomous TE expression from gene-derived signals [15]. These challenges are especially pronounced when attempting single-locus TE quantification, which requires assigning each read to the exact insertion from which it originates. Because this is rarely achievable with certainty, many studies aggregate TE expression at the family or subfamily level [15]. While computationally convenient, this approach obscures substantial within-family heterogeneity in which specific insertions rather than whole TE families drive regulatory activity [16]. The identity of the expressed locus is often critical: insertions at different genomic locations can carry distinct epigenetic states, regulatory potential, and effects on nearby genes [16, 17, 18]. Thus, resolving TE activity at single-locus resolution is essential for dissecting TE biology and its influence on gene regulation.

These challenges have motivated the development of several computational methods aiming to quantify TE expression from RNA-seq. Tools differ in their mapping strategies, treatment of multi-mapping reads and approaches for discriminating gene-derived from TE-derived signal [19, 20]. Most were originally developed for bulk RNA-seq. scRNA-seq data are sparser, have lower read depth per cell and often display strong 3’ or 5’ biases, all of which exarcebate ambiguities in locus-level TE quantification [21]. To date, five methods are explicitly designed for single-cell TE quantification [22, 18, 23, 24, 25], and only some of them [18, 23, 25] attempt locus-specific quantification.

Benchmarking efforts have so far been limited. Two independent studies evaluated bulk RNA-seq methods for locus-level TE quantification [19, 20], but these assessed different sets of tools and yielded conclusions that cannot be directly compared. This highlights the need for reproducible, extensible benchmarking pipelines that can incorporate newly developed tools [26]. Moreover, bulk benchmarks do not directly translate to single-cell settings, since they involve different challenges [27].

Existing single-cell benchmarks are smaller in scope and/or restricted to subfamily-level quantification. For example, the developers of SoloTE [18] compared their method only with scTE [22] using simulated short-read scRNA-seq data. Similarly, the IRescue study [24] benchmarked the method against both SoloTE and scTE and reported superior performance. However, these comparisons were performed exclusively at the TE subfamily level, since locus-level quantification is not supported by either IRescue or scTE. The only evaluation explicitly targeting locus-level TE quantification was presented in the publication introducing Stellarscope [23]. The authors compared the total number of unique molecular identifiers (UMIs) assigned to TE annotations across methods, but did not assess accuracy at individual loci. A more comprehensive evaluation was not possible because neither experimental nor simulated ground-truth data were available. Finally, MATES [25] was evaluated against scTE and SoloTE, but solely in terms of downstream cell clustering performance, again without the use of a known ground truth for TE expression. As a result, the performance of existing tools as well as the feasibility and limitations of locus-level TE quantification in scRNA-seq remain unclear.

To address this gap, we developed a comprehensive and reproducible benchmarking framework for evaluating locus-specific TE quantification methods using scRNA-seq data. We focused on short-read, droplet-based scRNA-seq, currently the most widely used modality [28] and the data type for which most single-cell TE quantification tools have been designed and evaluated. Our approach combined real datasets with a simulation pipeline that generated read-level ground truth, enabling direct and controlled evaluation of detection accuracy, quantification performance, and gene–TE disambiguation. We implemented a flexible Snakemake workflow [29] that standardizes preprocessing from raw sequencing reads, enabling multiple tools to be evaluated under consistent conditions. Using this framework, we systematically assessed how methodological choices, such as different strategies for handling multi-mapping reads and approaches for distinguishing TE-from gene-derived signal, affect locus-level TE quantification. We further analyzed how intrinsic properties of expressed TE loci, including evolutionary age and family origin, influence quantification performance.

The resulting resource is fully reproducible and readily extensible to new datasets and tools. Beyond providing practical guidance for users, our results highlight inherent, sequence-driven constraints of short-read scRNA-seq for analyzing repetitive elements and delineate the current limits of locus-specific TE analysis at single-cell resolution.

## 2 Results

### 2.1 Benchmarked tools

In our evaluation, we included two methods for single-cell TE quantification that enable locus-specific assignment for at least a subset of reads: SoloTE [18] and Stellarscope [23] (an adaptation of Telescope [30] for single-cell analysis; Table 1). In addition, we included STARsolo [31], a scRNA-seq preprocessing tool that performs read alignment, barcode demultiplexing, and gene quantification. Although not specifically designed for TE analysis, STARsolo supports joint gene and TE quantification and provides options for assigning multi-mapped reads to single loci. Its inclusion allowed us to assess how a general-purpose approach compares with TE-specific methods.

**Table 1:** Characteristics of TE locus-level quantification methods included in our benchmarking.

| Method | Multimappers included | Locus selection | Gene/TE distinction | Gene counts | Extra features | Year of publication |
| --- | --- | --- | --- | --- | --- | --- |
| STARsolo<sup>1</sup> | ✗ | Best aligned | ✓ | ✓ | – | 2021 |
| STARsolo<sup>1</sup> with EM | ✓ | EM | ✓ | ✓ | – | 2021 |
| SoloTE<sup>2</sup> | ✗ | Best aligned | ✓ | ✓ | Aggregated counts<sup>6</sup> | 2022 |
| SoloTE<sup>3</sup> with threshold | Above MAPQ thr.<sup>5</sup> | Best aligned | ✓ | ✓ | Aggregated counts<sup>6</sup> | 2025 |
| Stellarscope<sup>4</sup> | ✓ | EM | ✗ | ✗ | Cell pooling<sup>7</sup>, UMI deduplication | 2025 |

STARsolo operates directly on raw sequencing reads provided as FASTQ files. In contrast, SoloTE and Stellarscope require precomputed alignment files as input.

One key difference between these tools lies in their handling of multi-mapping reads. SoloTE selects the best alignment for each read, providing locus-specific quantification only for uniquely mapped reads (MAPQ score equal to 255). By default, multi-mapping reads are considered only at the subfamily-level. Here, for our benchmarking, we modified SoloTE to generate also locus-level counts for multi-mapping reads by adjusting the MAPQ threshold (see Sec. 5.5 and Table S1). Instead, Stellarscope uses an EM algorithm to iteratively reassign multi-mappers. This algorithm maximizes the posterior probability of a read mapping to a specific locus, calculated as the product of the proportion of total and multi-mapping UMIs mapping to that locus, and their mapping qualities. EM can be applied at the single-cell level or after pooling cells (e.g. by cell type or as a pseudobulk) to reduce sparsity. In both cases, the resulting UMI count matrices are calculated at the single-locus and single-cell levels. Unless otherwise specified, we consider Stellarscope in a pseudobulk mode. Similar to Stellarscope, STARsolo uses an EM algorithm with Maximum Likelihood Estimation to allocate multi-mapping UMIs, taking into account other UMIs (both uniquely and multi-mapped) from the same cell only.

The second key difference between the tools concerns how they distinguish reads originating from TEs versus genes. SoloTE first filters out reads that map entirely within exonic regions (which are assumed to originate from genes), before intersecting the remaining reads with annotated TE regions. This intersection is performed using bedtools [32] with default parameters, hence the strand which a read maps to is not taken into account. SoloTE then generates count matrices that include expression values for both genes and TEs. Stellarscope, instead, performs the intersection with the TE annotation directly, without excluding any gene-mapped reads beforehand, using the intervaltree [33] Python package, taking into account the strandeness. A similar approach can also be enabled in SoloTE using the “dual” option. In our implementation of STARsolo for TE quantification, reads are mapped to all features present in the provided annotation, which included both gene transcripts and TE loci.

A unique feature of Stellarscope is its multimapper-aware UMI deduplication step. Rather than resolving multimapping to a single locus prior to UMI deduplication, Stellarscope collapses reads that share a UMI and any overlapping set of candidate loci. Reads with the same UMI but mapped to different sets of loci are instead counted separately, as these are likely derived from distinct molecules that coincidentally received the same UMI.

### 2.2 Locus-specific TE transcription in real scRNA-seq data: patterns and methodological comparisons

#### 2.2.1 Abundant and informative locus-level TE signal in single-cell transcriptomes

First, we aimed to establish how much locus-level TE signal is present in real scRNA-seq data. We analyzed three publicly available datasets and characterized the abundance, mapping properties and biological relevance of TE-derived reads at single-locus resolution. All datasets generated using 10X Genomics platforms (see Sec. 5.1). The first consisted of mouse embryonic stem cells (mESCs) differentiating into 2-cell-like-cells (2CLC) [34], a totipotent-like cell state known to exhibit widespread upregulation of multiple TE families [2, 35, 36]. The second was derived from the mouse olfactory mucosa [37], a tissue characterized by continuous cell turnover and active chromatin remodelling [38, 39], making it a relevant context in which to explore TE transcriptional activity. The third is a human peripheral blood mononuclear cell (PBMC) dataset obtained from the 10XGenomics database [40], comprising multiple well-characterized cell types and widely used as a benchmark for evaluating scRNA-seq analysis methods [41].

As an initial analysis, we quantified the fraction of reads mapping to TE loci compared to gene regions. For this purpose, we used SoloTE with default parameters, which performs direct read assignment without EM-based redistribution of multi-mapped reads (Sec. 2.1). Across all datasets, we observed that a substantial fraction of reads mapped to TE loci, exceeding 24% in every case (Fig. 1A). In standard scRNA-seq analysis pipelines, these reads would typically be discarded, either because they map to multiple genomic locations or because they fall outside conventional gene annotations, which generally exclude TEs. Among reads mapping to TEs, the proportion of multi-mapping reads varied across datasets (Fig. 1B). The mESC dataset containing 2CLCs showed the highest fraction of multi-mappers, consistent with the activation of evolutionarily young and highly repetitive TEs in this cell state (see Sec. 2.3.2). In contrast, somatic tissues such as PBMCs and olfactory mucosa exhibit a lower, though still substantial, proportion of multi-mapped TE reads.

**Figure 1:**
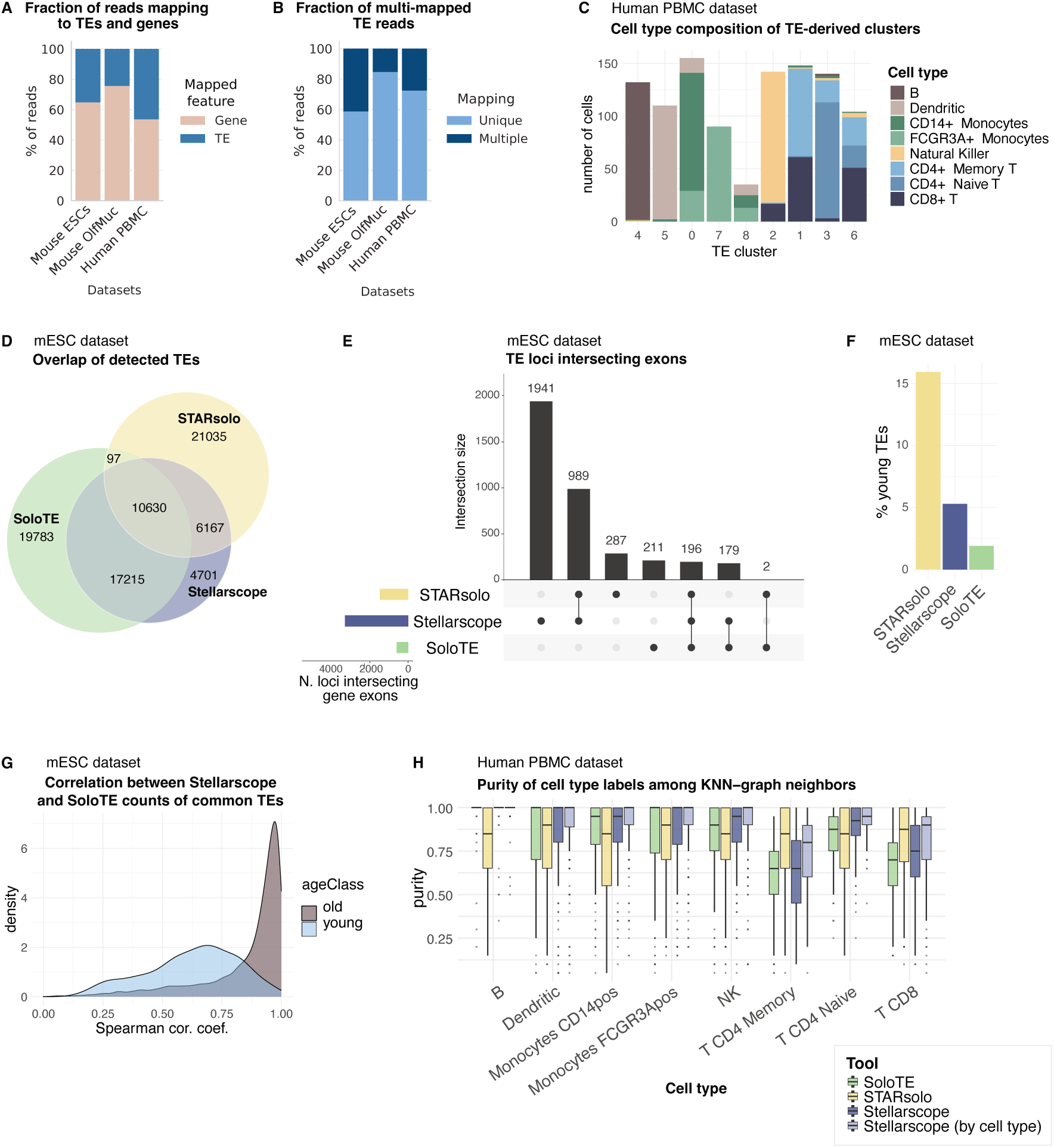
**A** Percentages of reads assigned to genes and TEs by SoloTE in each scRNA-seq dataset. **B** Percentages of reads uniquely mapped and multi-mapping among reads assigned to TEs by SoloTE. **C** Number of cells assigned to each gene-derived cell type within each TE-derived cluster in the human PBMC dataset. **D** Euler diagram of TE loci detected in the mESC dataset by SoloTE, STARsolo (with EM) and Stellarscope. **E** Upset plot of detected TE loci intersecting gene exons found by each tool in the mESC dataset. **F** Percentage of young TEs among those detected in the mESC dataset by each tool. **G** Distribution of Spearman correlation coefficients between normalized counts obtained with SoloTE and Stellarscope, calculated per locus across all cells. Only loci detected by both tools and STARsolo are included. **H** Boxplots of purity of cell type labels among the 20 nearest neighbors of each cell in the KNN-graph obtained with TE locus counts estimated by SoloTE, STARsolo with EM, Stellarscope run in pseudobulk mode and in celltype mode. Unless otherwise specified, Stellarscope was run in a pseudobulk mode.

To determine whether the TE signal captured at single-locus resolution reflects biologically meaningful structure, we next assessed whether TE-derived expression profiles could resolve cellular heterogeneity within each dataset. We analyzed gene and TE counts separately and performed cell clustering, selecting the resolution that maximized concordance between gene- and TE-based clusters using the Adjusted Rand Index (ARI). Across all analyzed datasets, major cell types were distinguishable based solely on TE locus–level counts (Supp. Fig. S1A–E). We focus here on the PBMC dataset, which contains the largest number of well-annotated and transcriptionally distinct cell types. Clustering based on TE profiles showed strong agreement with gene expression–derived cell type annotations (Fig. 1C). Minor discrepancies were primarily limited to the separation of closely related immune subtypes, such as CD14+ versus FCGR3A+ monocytes and among T cell subpopulations.

TE-based clustering identified different substructure that were not apparent in gene expression–derived clusters. For example, TE cluster 8 in the PBMC dataset (Fig. S1F) and TE-derived cluster 5 within pluripotent cells in the mESC dataset (Fig. S1G) were not resolved by gene-based analyses. Further investigation will be required to determine whether this substructure reflects biologically meaningful cell states or instead arise from technical or analytical factors.

Taken together, these findings demonstrate that: (i) there is a substantial amount of TE signal in scRNA-seq datasets, (ii) a large proportion of TE-associated reads are multi-mapped and therefore require careful handling, and (iii) even when restricting attention to uniquely mapped reads, TE expression alone is sufficient to resolve complex cellular compositions, potentially revealing biologically relevant subtypes or states that would be obscured when solely focusing on genes. These observations are consistent with previous reports [18, 23, 16], demonstrating cell type-and state-specific TE expression patterns in single-cell data.

#### 2.2.2 Comparison of TE quantification methods on real data

We next compared locus-level outputs of different TE quantification methods on the mESC and human PBMC datasets, where TE-mapped reads are more abundant. Specifically, we contrasted the SoloTE results presented above with those obtained using Stellarscope (pseudobulk mode) and STARsolo with EM-based quantification.

Focusing on the dataset with the highest fraction of TE-mapped reads (according to SoloTE), i.e., the mESC dataset, we first examined which TE loci were detected (above 1 count) by each tool in at least 2% of cells. Our analysis revealed a limited overlap across methods (Fig. 1D; a similar pattern was observed for the PBMC dataset, Fig. S2A). STARsolo exhibited the highest divergence with respect to the other tools, while SoloTE identified the largest number of TE loci overall (Fig. S2B). The discrepancy in number of loci between the different tools decreased when limiting the analysis to TEs detected in a higher percentage of cells (Fig. S2C), indicating that the additional loci detected by SoloTE are expressed in a small number of cells, possibly due to stochastic noise or higher cell specificity. To better understand the sources of these discrepancies, we next examined key features of the loci detected by each method, focusing on their genomic context and evolutionary age.

One potential concern in TE quantification is the misattribution of reads originating from gene transcripts to TE loci. This can occur when sequencing reads derive from exonic or intronic regions of genes that overlap annotated TEs, or when fragments from gene-derived transcripts are ambiguously assigned to repetitive TE sequences. In such cases, TE counts may reflect gene expression rather than autonomous TE transcription. To assess this effect, we analyzed the overlap between detected TE loci and annotated gene exons (Fig. 1E, Fig. S2D). For the mESC dataset, nearly half of the TE loci uniquely identified by Stellarscope overlapped gene exons (*n* = 1941 out of 4701). In contrast, SoloTE produced the lowest overlap. This difference is due to SoloTE’s filtering, which explicitly excludes reads mapping to gene annotations. While some overlapping loci may indeed represent genuine autonomous TE transcription within gene regions, not excluding fragments mapping to gene bodies (as in Stellarscope) may increase the risk of assigning gene-derived counts to TEs rather than to their true source transcripts.

We next stratified TE loci according to their evolutionary age. TE age is a critical factor because the nucleotide sequence of younger elements tends to be more similar to other members of the same family and thus more difficult to resolve at single-locus resolution. TEs were classified as “young” or “old” based on whether their estimated age was below or above 2 million years, respectively (Sec. 5.2). STARsolo detected the highest proportion of young TEs, followed by Stellarscope (Fig. 1F, Fig. S2E). This pattern may reflect increased sensitivity to young, repetitive elements due to the inclusion of multi-mapped reads and EM-based assignment strategies, but it may also indicate a higher false-positive rate for these loci.

For the mESC dataset, we next examined expression levels for TE loci that were consistently detected with at least one count in at least 2% of cells by all methods (*n* = 10630). For these loci, TE counts produced by the different methods were highly correlated for older TEs, whereas the correlation was markedly reduced for young elements across all datasets (Fig. 1G, Fig. S2F-J). This result is consistent with the higher repetitiveness of young TEs, making locus-level quantification particularly challenging.

Finally, we compared the methods in terms of their ability to capture cell type structure identified from gene expression. Using the PBMC dataset, we quantified the purity of cell type annotations within local neighborhoods of the k-nearest neighbor (KNN) graphs constructed from TE counts (Fig. 1H). Most cell types exhibited high annotation purity across TE quantification methods. However, STARsolo consistently produced lower purity scores (except for CD4+ T Memory cells and CD8+ T cells). SoloTE and Stellarscope in pseudobulk mode showed similar performance, with Stellarscope achieving equal or slightly higher purity in all cell types. Providing cell type labels to Stellarscope and fitting the EM model separately for each cell type further improved annotation purity. The increased similarity to gene-based annotations obtained by Stellarscope may reflect improved capture of cell type-specific TE expression, but it could also arise from the potential misattribution of gene-derived reads to TEs (Fig. 1E).

In summary, our analysis revealed substantial and biologically structured TE expression at single-locus resolution, while also highlighting pronounced differences among TE quantification methods. However, the absence of a ground truth precludes a rigorous assessment of which approach most accurately reflects true TE transcriptional activity. To address this limitation, we next benchmarked the methods using simulated data with known TE expression levels.

### 2.3 Benchmarks using simulated ground truth

We devised a simulation framework that enables quantitative evaluation of each tool’s performance by generating scRNA-seq reads from user-defined TE count matrices. This design allows direct comparison between inferred and true locus-level TE counts. In this section, after describing the generation of these synthetic datasets, we use them to assess each tool’s ability to detect TE loci, accurately quantify their expression, and resolve gene-TE assignment ambiguity under controlled conditions.

#### 2.3.1 Synthetic data generation

The overall simulation workflow is schematically depicted in Fig. 2A (see also Sec. 5.3).

**Figure 2:**
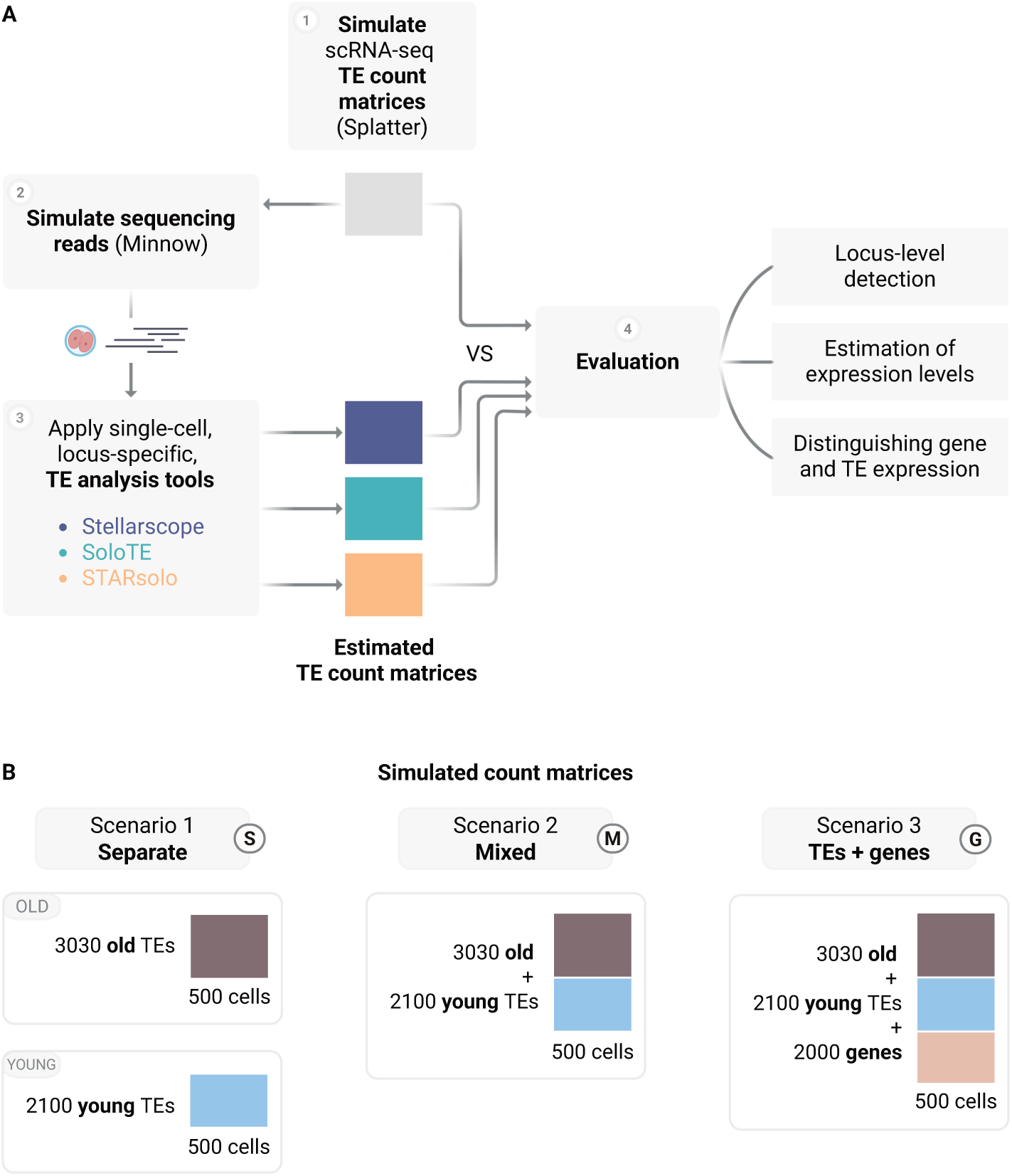
**A** Schematic of our benchmarking workflow. **B** Simulated count matrices included in the benchmarking.

In brief, we used Splatter [42] to simulate scRNA-seq TE count matrices. We generated balanced representations of TE families in mouse and ensured that both young and old loci were simulated at comparable expression levels. Although this may not recapitulate real TE expression patterns, this design allowed us to disentangle effects driven by expression magnitude from those driven by sequence repetitiveness or evolutionary age. We generated multiple configurations of increasing complexity, so that the causes of each tool’s performance could be precisely traced (Fig. 2B). In total, we produced three TE-only matrices: one containing exclusively old TEs (Scenario 1, “Separate old”), one exclusively young TEs (Scenario 1, “Separate young”) and one containing both (Scenario 2, “Mixed”). Furthermore, to assess how gene expression interferes with TE quantification, we also generated a fourth matrix containing both young and old TEs together with 2,000 highly variable genes derived from the mESC dataset (Scenario 3, “TEs+genes”, Fig. 2B). To convert these simulated TE count matrices into realistic droplet-based scRNA-seq reads, we used Minnow [43]. Minnow models several characteristics of droplet-based scRNA-seq, including: multimapping behaviour (combining inherent sequence-level ambiguity with the chosen read length and feature-level ambiguity learned from real data), PCR amplification biases, UMI and sequencing errors and fragmentation patterns. Simulations were trained on the mESC dataset to mimic its multi-mapping rates.

Starting from these simulations, we evaluated each tool under four aspects: (i) ability to detect which TE loci are expressed in each cell (Sec. 2.3.2); (ii) accuracy in estimating expression levels (Sec. 2.3.3); (iii) performance at the TE-family level (Sec. 2.3.4); (iv) ability to distinguish TE-derived from gene-derived reads (Sec. 2.3.5).

#### 2.3.2 Detection of expressed TEs

First, we assessed the ability of different tools to correctly identify expressed TE loci, focusing on differences between young and old elements across the three TE-only simulation settings. The fraction of multi-mapping reads in the mixed simulation (Scenario 2) is within the range observed in the real datasets (Fig. 3A). This indicates that our simulation reproduced realistic mapping ambiguity and provided an ideal testing ground for evaluating tool performance under controlled conditions. As it may be expected, older TE loci predominantly generated uniquely-mapping reads, while over 80% of reads derived from younger loci mapped to multiple locations (Fig. 3A). The multi-mapability issue of TE sequencing data [15] therefore mainly affects young insertions. This also suggests that the higher rate of multi-mappers found in the mESC dataset compared to other samples (Fig. 1A) reflects more abundant expression of young TEs in 2CLCs.

**Figure 3:**
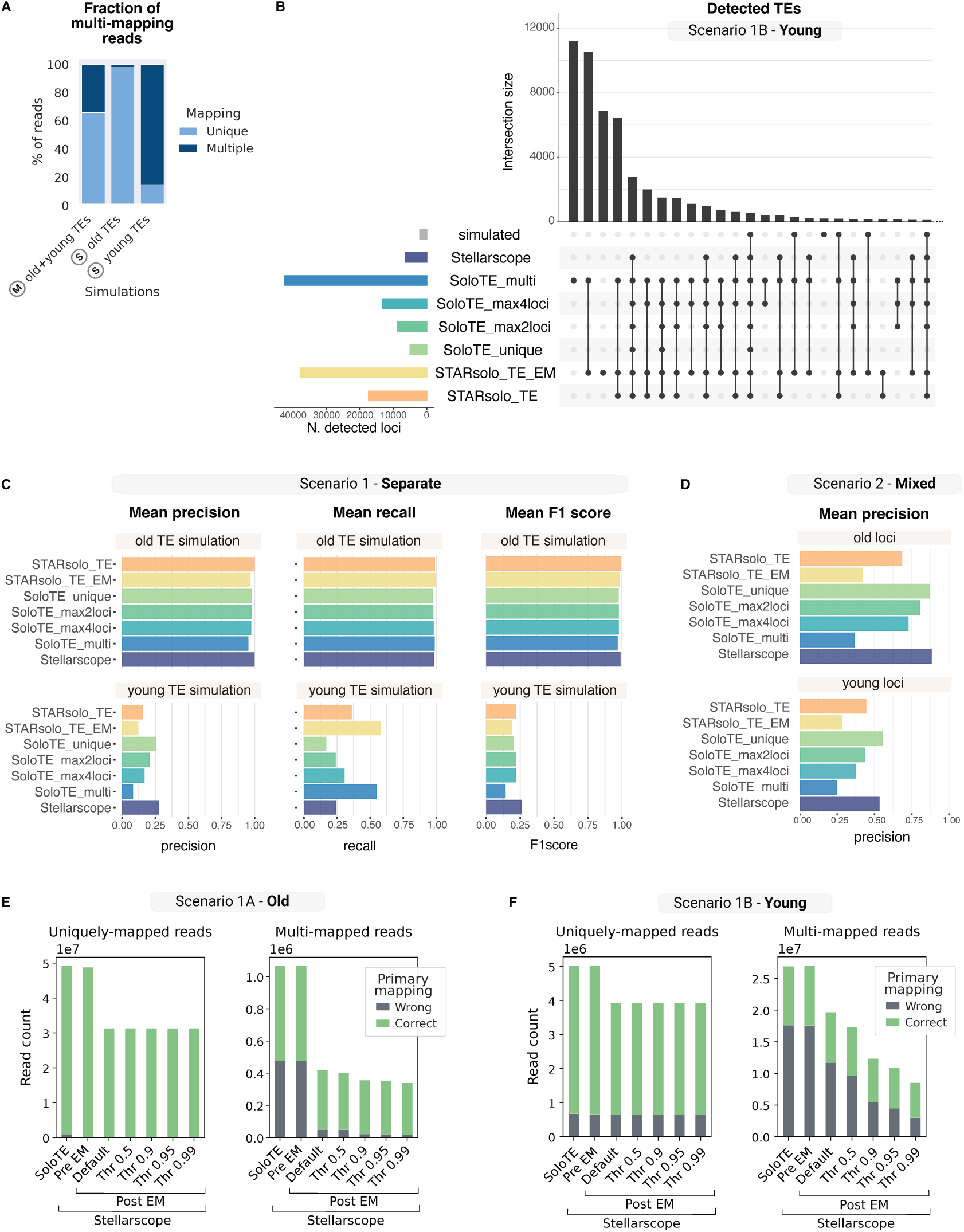
**A** Percentage of uniquely and multi-mapping reads in each simulation (derived from SoloTE). **B** Upset plot of sets of TE loci present in the young TE simulated matrix (grey bar) and detected by each tool. SoloTE settings listed in Table S1. **C** Precision, recall and F1 score computed per cell and then averaged across cells, using binarized TE count matrices. Results are shown for simulations with old loci (top row) and with young loci (bottom row). **D** Precision computed per cell and then averaged across cells, using the binarized count matrix obtained from the simulation containing both old and young loci. Results are shown separately for old and young loci. **E,F** Number of primary alignments correctly and incorrectly assigned to their true originating TE locus in simulations containing only old (E) or only young (F) TE insertions. Reads are stratified by mapping uniqueness: uniquely mapped reads (MAPQ = 255) versus multi-mapped reads (MAPQ lower than 255). For Stellarscope, results are shown pre-EM (initial alignment-based assignment), post-EM default mode (i.e., best exclude mode, reads assigned to the feature with highest posterior probability, excluding ties), and post-EM with varying posterior probability thresholds (0.5, 0.9, 0.95, 0.99)

Leveraging the ground truth, we quantified the number of loci detected by each tool across all cells, treating detection as a binary event (i.e., any non-zero count). When simulating old TE loci alone (Scenario 1 old), the number of detected loci closely matched the ground truth across all methods (Fig. S3A). In contrast, in the young TE simulation (Scenario 1 young), loci were consistently over-detected, with all tools reporting a substantial amount of false positives (FPs), with limited agreement across tools (Fig. 3B). Tools that incorporated multi-mapped reads (Stellarscope, STARsolo with EM, and SoloTE with multi-mappers) exhibited a larger FP burden than SoloTE restricted to uniquely mapped reads. Among these, Stellarscope was the most conservative, suggesting that its EM algorithm partially mitigates the noise introduced by multi-mapping.

In Scenario 1 young, 2,768 FP loci were detected by all tools (common FP), whereas 193 simulated loci were not detected by any method (common false negatives; FN), suggesting shared biases inherent to locus-level TE quantification. Family composition differed markedly between simulated loci, common FPs and common FNs (Fig. S3B; *χ*^2^ = 860, *df* = 6, *p <* 2 × 10*^—^*^16^). L1 elements were enriched among the common FNs, indicating systematic under-detection. In contrast, the common FP loci were enriched for LTR elements (ERVL, ERVK, ERV1), were significantly longer (Fig. S3C; Wilcoxon rank-sum test, *p <* 2 × 10*^—^*^16^) and had higher sequence divergence (Fig. S3D; Wilcoxon rank-sum test, *p <* 2 × 10*^—^*^16^) than the simulated loci, consistent with misassignment of reads derived from young loci to older elements.

We next computed precision and recall per cell (Fig. 3C). In the simulation with only old TEs (Scenario 1 old), both metrics were consistently high, with STARsolo (without EM) and Stellarscope offering a slightly higher F1 score (harmonic mean of precision and recall). In the young TE simulation (Scenario 1 young), detection was uniformly poor: even the best-performing method (Stellarscope) achieved an F1 score below 0.3. When analysing the simulation containing both young and old TEs (Scenario 2, the closest to real data), recall remained comparable to that of Scenario 1 old (Fig. S3E). Precision for old loci dropped, whilst it increased for young loci (Fig. 3D). This pattern appears because reads originating from young elements were partially misassigned to older loci, even by tools restricted to uniquely mapped reads (e.g., SoloTE). Thus, in real datasets containing TEs of all ages, counts for old loci may be indirectly affected by mapping ambiguity, highlighting that relying solely on unique mappers does not fully bypass multi-mapping artifacts. In the mixed simulation, Stellarscope and SoloTE with uniquely mapping reads outperformed the other tools in the detection of old loci. STARsolo (both with and without EM) instead achieved the highest F1 score for young TEs, driven by a higher recall. For young elements in the mixed simulation (Scenario 2), SoloTE with all multi-mappers included surpassed both the other SoloTE configurations and Stellarscope (Fig. S3C-D). In the mixed simulation, Stellarscope and SoloTE restricted to uniquely mapping reads outperformed the other tools in detecting old loci. In contrast, for young TEs, STARsolo (with and without EM) achieved the highest F1 score, and SoloTE configured to include all multi-mapping reads outperformed all the other SoloTE configurations and Stellarscope, primarily driven by higher recall (Fig. S3C–D).

To evaluate whether filtering lowly expressed loci improves detection performance, we computed precision–recall (PR) curves by varying the minimum count threshold required to classify a locus as detected (1–100 counts). For older loci, increasing the threshold up to 10 counts per cell modestly improved precision for most methods, particularly those with higher false-positive rates at the lowest threshold (Fig. S4A). STARsolo (without EM) and Stellarscope consistently achieved the highest precision across thresholds. Due to the simultaneous drop in sensitivity, however, the best F1 score is obtained keeping a threshold of 1 (Fig. S4B). Stricter filtering is therefore not recommended for older elements. Correspondingly, the area under the PR curve (AUPRC) remained high for the old simulation, indicating that detection of older TEs is robust to the choice of count threshold (Fig. S4C). Performance for young TEs was substantially poorer based on AUPRC. Precision increased rapidly with higher thresholds, but recall (already low at baseline) dropped, reflecting a pronounced sensitivity-specificity trade-off (Fig. S4D). Stellarscope achieved the highest F1 score at the lowest threshold, whereas STARsolo and SoloTE (multi-mapper mode) outperformed other methods under more stringent filtering (Fig. S4E). Overall, in the mixed simulation (Scenario 2, Fig. S4F), the 1 count threshold remained the one resulting in the best F1 score. These results demonstrate that count-based filtering primarily shifts the balance between sensitivity and specificity rather than uniformly improving performance, and that the magnitude of this trade-off depends strongly on TE age and multi-mapping behavior. Accordingly, these PR curves should be interpreted as method-specific performance profiles rather than universal recommendations for expression thresholds, as optimal cutoffs will vary across datasets and expression patterns.

Given the high FP rates observed for young TEs across tools, we further investigated the underlying causes and potential mitigation strategies by assessing locus-level assignment accuracy at the single-read level. We focused on SoloTE and Stellarscope, as both methods give in output modified alignment files that report per-read feature assignments. In particular, Stellarscope provides assignments both before and after EM, enabling an explicit evaluation of the impact of its probabilistic read reallocation. It also reports a locus-specific posterior probability for each read, interpreted as a confidence score, which can be used to filter the resulting count matrix (by default, reads assigned with equal probabilities to two or more loci are excluded). We applied increasingly stringent thresholds to assess whether higher confidence filtering improved assignment accuracy. Consistent with our earlier results, reads originating from old TEs (Scenario 1 old; Fig. 3E) were assigned with near-perfect accuracy when uniquely mapped, even without EM (left panel). Activating the EM algorithm markedly reduced the number of incorrectly assigned reads (right panel), reflecting more conservative assignment. However, this increased stringency also resulted in the exclusion of a subset of correctly assigned reads from the final TE counts. A similar pattern was observed for young TEs (Scenario 1 young; Fig. 3F), although overall accuracy was considerably lower. Even uniquely mapped reads from young loci were frequently misassigned, likely due to simulated sequencing errors that promoted erroneous alignments to highly similar loci. Across both scenarios, Stellarscope assignments without EM were broadly comparable to those produced by SoloTE. EM improved assignment accuracy by favoring higher-confidence allocations, but at the cost of discarding a substantial fraction of correctly assigned reads, particularly for older loci. Applying an additional posterior probability filters further increased stringency. This tradeoff was reflected in the locus-level detection accuracy of Stellarscope after applying a posterior probability filter of 0.9, which resulted in slightly higher precision but lower recall compared to the default settings, while the overall F1 score remained similar (Fig. S5A).

Taken together, these results show that locus-specific detection of old TEs is generally accurate across methods, whereas young, more repetitive elements remain difficult to quantify and are consistently over-detected. Multi-mapping is the primary source of error, affecting both EM-based and unique-mapper strategies, and even influencing quantification of older loci when young elements are present. EM and posterior filtering increase precision but reduce sensitivity, highlighting persistent trade-offs.

#### 2.3.3 Estimation of expression levels

We next assessed how different methods compare in terms of per-cell TE expression levels for the loci they identify. We restricted this analysis to true positives (TPs), i.e., loci that were both simulated and correctly detected, thus isolating expression-level accuracy from detection performance.

To quantify agreement between simulated and estimated expression values, we computed the Spearman’s rank correlation for each method, separately for the simulations with only old or young TEs (Scenario 1). Results differed markedly between age groups (Fig. 4A). For old TEs, estimated counts closely matched the simulated expression levels (Fig. 4B, S6A), with all methods achieving a correlation close to 1. By contrast, quantification of young TEs was substantially noisier (Fig. 4C, S6B): rank correlations dropped considerably for all tools, reflecting the inherent difficulty of allocating reads derived from highly similar loci. STARsolo showed modestly higher correlations, though differences between tools were small relative to the large overall gap between old and young TEs.

**Figure 4:**
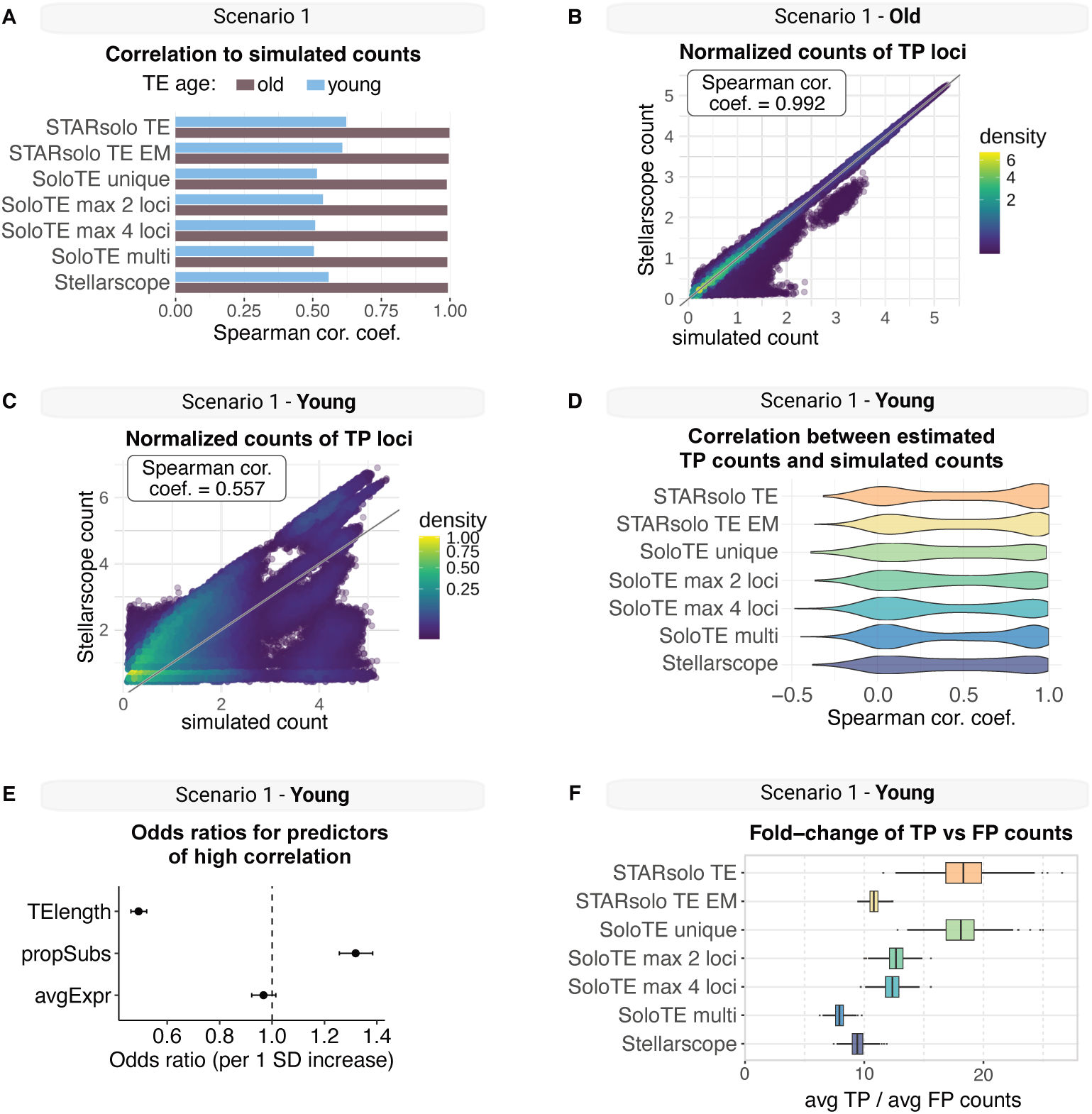
**A** Spearman correlation coefficients between true positive counts from each tool and simulated counts, calculated separately for old and young loci. **B, C** Scatterplot of normalized simulated counts (x-axis) versus Stellarscope counts (y-axis) for correctly detected old (B) and young (C) loci. Points are colored by local density, estimated using two-dimensional kernel density estimation (kde2d, MASS package [44]) with bandwidths determined by Silverman’s rule of thumb [45]. **D** Distribution of Spearman correlation coefficients between estimated and simulated counts in the young TE simulation. Correlations were computed separately for each correctly detected locus and for each tool. **E** Odds ratios (points) and 95% confidence intervals (horizontal bars) from a mixed-effects logistic regression model predicting whether a TE locus exhibits high correlation accuracy (above-median correlation). Predictors were standardized (mean 0, SD 1) so effect sizes represent the change in odds per one standard deviation increase. **F** Per-cell ratio of average raw counts for TP versus FP young loci, by tool.

When correlations were computed on a per-locus basis rather than pooled across loci, we observed a large spread across loci, with a bimodal pattern (Fig. 4D). Given the high spread, we investigated the causes behind the different quantification accuracy of different loci. To identify locus-level features associated with higher agreement to ground truth, we fitted a mixed-effects logistic regression model with high versus low correlation (defined relative to the median correlation across loci) as a binary outcome and tool included as a random intercept to account for baseline performance differences among tools (see Sec. 5.7). After accounting for between-tool variability (random intercept variance = 0.025), both TE length and sequence divergence were significant predictors of accuracy (Fig. 4E). Specifically, longer TEs were less likely to exhibit high correlation (*β* = —0.71 *±* 0.03 SE, *p <* 2 × 10*^—^*^16^). In contrast, higher sequence divergence relative to the consensus (corresponding to relatively older elements within the young TE class) was associated with increased accuracy (*β* = 0.28 *±* 0.02 SE, *p <* 2 × 10*^—^*^16^). Average ground truth expression was not significantly associated with accuracy after adjusting for other predictors (*β* = —0.03 *±* 0.02 SE, *p* = 0.17).

Finally, we examined the expression levels assigned to FP young loci, i.e., loci incorrectly reported as expressed, to determine whether they were at least quantified at lower levels than true positives. Indeed, FP loci showed substantially lower average expression than TPs (Fig. 4F, S6C), with the clearest separation observed in methods that rely exclusively on unique mappers (STARsolo-TE and SoloTE-unique). This pattern indicates that FP loci often arise from low-level noise or sparse multi-mapping events. Expression levels of loci that were not detected (FNs) were also generally lower than those of TPs (Fig. S6D), although the difference between the two was less pronounced than in the case of FPs. The effect of filtering low-abundance features in the young TE simulation was shown in Sec. 2.3.2.

In summary, expression-level quantification was highly accurate for older TEs across all tools, but substantially degraded for younger elements, whose repetitive nature introduces pervasive assignment noise.

#### 2.3.4 Family-level analysis

The challenges observed in quantifying young TEs at single-locus resolution raise the question of whether TE family membership influences quantification accuracy. Families differ widely in their evolutionary age, copy number, length and sequence divergence, leading to substantial heterogeneity in mappability and read ambiguity. Consequently, the ability of computational methods to correctly detect and assign reads to individual loci may depend strongly on the family to which those loci belong. Understanding these family-specific effects is essential, as biological systems often display selective activation of particular TE classes, and mis-quantification at the family level can bias downstream interpretations of TE activity.

We therefore repeated the assignment-accuracy analysis described in Fig. 3C, restricting it to the young TE simulation and stratifying them by TE family. This revealed pronounced variation in performance across families (Fig. 5A). Among the families examined, L1 elements exhibited the poorest combination of precision and recall across all tools, followed by the ERVL family. By contrast, Alu elements showed the highest precision, with both SoloTE and Stellarscope achieving values above 0.7 and a higher F1 score compared to other families. B2 elements, the other SINE family included in our simulations, ranked second. These differences may reflect fundamental sequence properties: SINEs are shorter (Fig. S7A) and likely to display greater sequence divergence within read-covered regions (Fig. S7B), reducing multi-mapping ambiguity relative to the longer, more homogeneous L1 and ERVL elements. Tool performance also varied in a family-specific manner. Solo-TE (unique-mapper mode) achieved substantially higher precision than other tools for L1 elements, whereas Stellarscope outperformed SoloTE for ERV1 elements.

**Figure 5:**
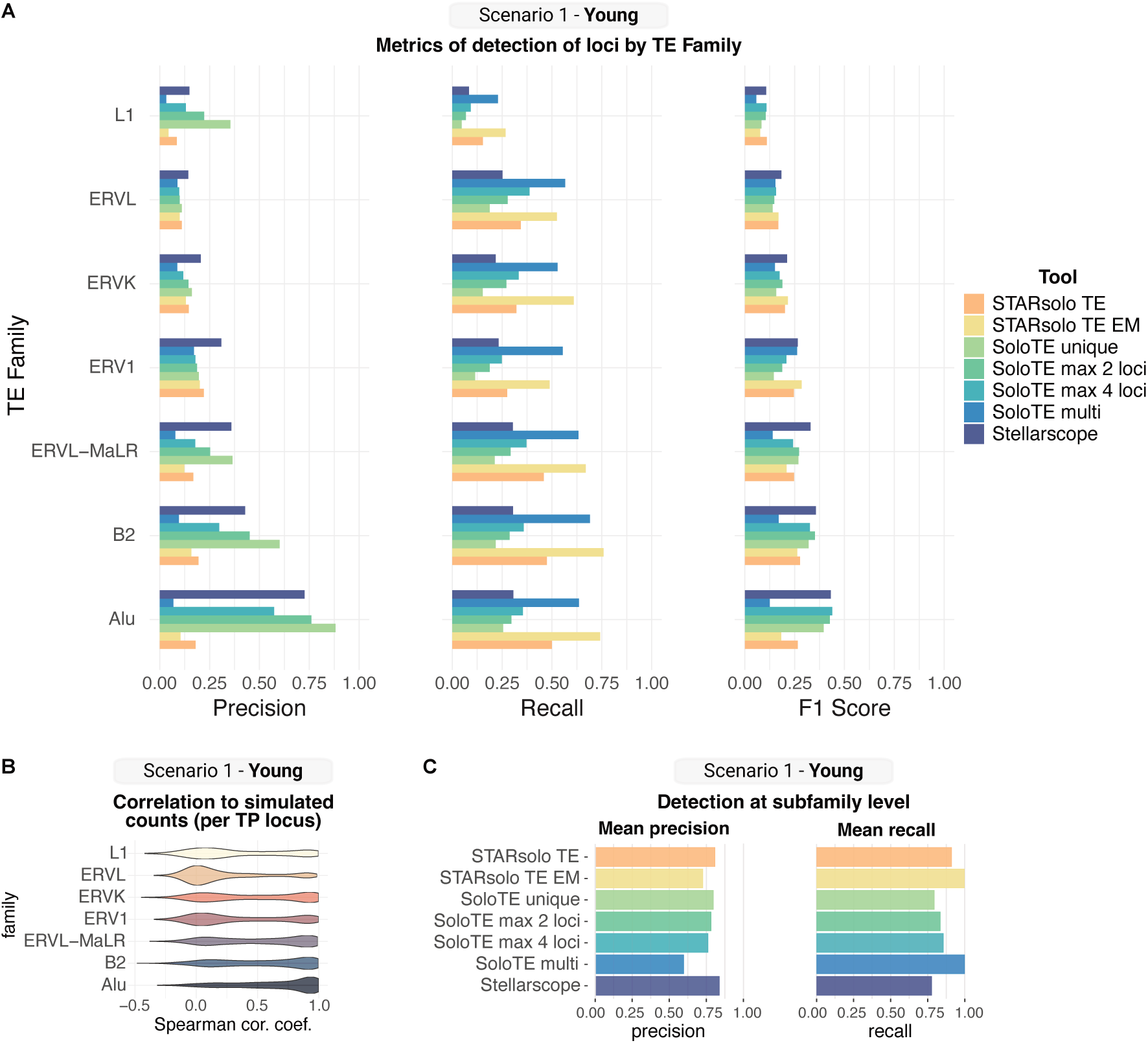
**A** Precision, recall and F1-score computed separately for each TE family in the young TE simulation; metrics were computed per cell and then averaged across cells. **B** Spearman correlation between true positive loci counts and simulated counts (Scenario 1, young simulation), computed by TE family. **C** Precision and recall computed per cell and then averaged across cells, using binarized, subfamily-level aggregated matrices from the young TE simulation.

Accuracy in estimation of expression levels also varied by family, with ERVL elements showing the largest portion of low correlations to the ground truth (Fig. 5B). These contrasting patterns highlight that no single method performs optimally for all TE families. Instead, optimal tool selection may depend on the TE composition of the biological system under study, particularly in contexts where specific families or subfamilies are known to be transcriptionally more active.

Given the consistently low performance observed for young elements at the locus level, we next asked whether these elements could at least be accurately quantified at the subfamily level. To this end, we aggregated locus-level counts from each tool by subfamily and computed detection metrics against a correspondingly aggregated ground-truth matrix. Although some false positives remained (Fig. S7C) aggregation markedly improved precision across all methods compared to locus-specific quantification (Fig. 5C), indicating that reads were generally assigned to the correct subfamily even when misallocated among individual loci. Recall at the subfamily level also increased, suggesting that most expressed subfamilies were successfully detected. Further aggregation to the family level led to additional gains in precision and yielded near-perfect recall for all tools (Fig. S7D). Collectively, these results confirm that the core difficulty in TE quantification lies not in detecting TE expression *per se*, but in resolving expression at the level of individual, highly similar loci within a family.

#### 2.3.5 Evaluating cross-assignment between genes and TE loci

The simulated dataset containing both genes and TEs (Scenario 3, see Fig.2B) enabled us to examine a key source of error in locus-level TE quantification: the misassignment of reads between gene models and TE annotations. This issue is particularly important because TE insertions frequently overlap gene bodies, especially within introns and untranslated regions [15]. In real scRNA-seq data dominated by short reads and sparse coverage, such overlaps make it difficult to determine whether a read derives from an autonomous TE transcript, a gene-derived RNA that happens to contain TE sequence, or an ambiguous fragment that aligns with both. Understanding these sources of error is essential, as incorrect redistribution of reads can either inflate apparent TE activity or mask genuine TE-driven transcriptional signals.

We found that all methods detected expressed TE loci that overlapped annotated gene bodies (Fig. 6A). Some of these were true positives, i.e., loci simulated as expressed in gene-overlapping regions. However, the majority represented false positives, indicating incorrect allocation of gene-derived reads to TE loci. SoloTE, when restricted to uniquely mapping reads, produced the fewest false positives, whereas methods incorporating multi-mappers detected many more gene-overlapping TE loci. To dissect this further, we examined whether these gene-overlapping TE calls arose from genes that were actually expressed in the simulated matrix. The red bars in Fig. 6A mark the number of TE loci overlapping expressed genes, a sign of systematic misattribution of gene-derived reads to TE annotations. All methods showed some degree of this error, but Stellarscope exhibited the most pronounced effect, with more than 400 detected TE loci corresponding to expressed genes. This behavior reflects Stellarscope’s design choice of not filtering out reads mapping within gene bodies. Although one potential mitigation strategy is to remove gene-overlapping TE loci from the TE annotation, such filtering would preclude detection of genuinely expressed TEs located inside or near genes, which is an important class of regulator insertions.

**Figure 6:**
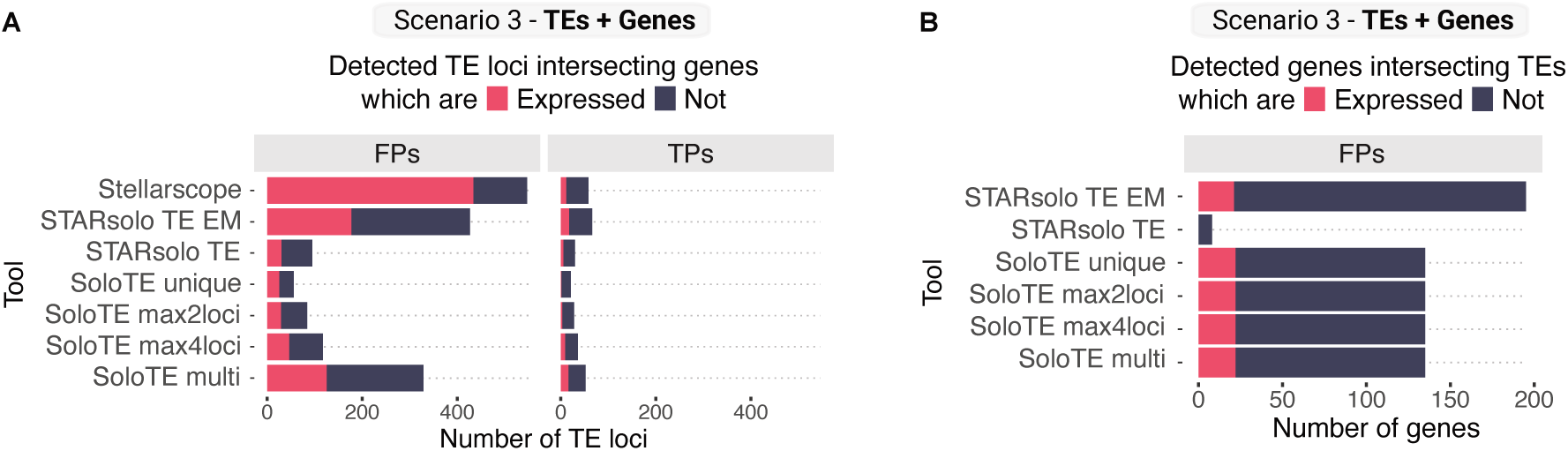
**A** Number of TE loci detected by each tool which overlap gene exons. TP (detected loci that were present in the simulated ground truth matrix) and FP (detected loci not present in the simulation) are shown separately. The red portion of the barplot shows the number of loci for which the overlapping gene is expressed. **B** Number of genes erroneously detected by each tool which overlap TE loci. The red portion of the barplot shows the number of genes for which the overlapping TE insertion is expressed.

The reverse process, i.e., misassigning TE-derived reads to genes, also occurred. In the mixed simulation (Scenario 3, TEs + genes), both SoloTE and STARsolo with EM falsely identified expression of genes overlapping expressed TEs (Fig. 6B). Surprisingly, this issue persisted even in the young-TE-only simulations, where no gene expression was simulated: SoloTE incorrectly called more than 200 genes as expressed based on TE-derived reads alone.

Together, these analyses show that gene-TE disambiguation remains a major challenge for locus-level TE quantification. Errors arise in both directions (genes misclassified as TEs and viceversa) and tools differ in how they prioritize gene versus TE models during assignment. Accurate separation of gene- and TE-derived reads will therefore require improved annotations, better modeling of shared sequences or hybrid strategies that explicitly incorporate gene and TE structure during alignment and quantification.

## 3 Discussion

In this study, we present a systematic benchmark of computational tools for quantifying TE expression at single-locus resolution from short-read scRNA-seq data. By combining analyses of real datasets with simulations that provide a ground truth, our work clarifies both the capabilities and the intrinsic limitations of current methods when applied to droplet-based single-cell sequencing assays.

Across real datasets, we observed that TE-derived reads constitute a substantial fraction of the transcriptome and capture biologically meaningful structure, yet their reliable quantification is hampered by widespread multi-mapping, particularly for evolutionarily young elements. Our simulations were designed to disentangle ambiguity caused by expression level, TE age, and genomic context. Our analyses revealed that locus-level quantification is highly accurate for older TEs, whereas quantification of young elements remains challenging for all tools, despite their attempts to resolve multi-mapped reads.

Among the evaluated methods, SoloTE (default mode, i.e., unique mappers only) and Stellarscope (with EM-based reallocation) showed broadly comparable performance across most tasks. Allowing multi-mappers in SoloTE substantially increased false positives without improving locus-level accuracy, and we therefore discourage this configuration. The use of EM in Stellarscope outperformed STARsolo, likely because Stellarscope pools information across cells (pseudobulk or cell-type modes), mitigating sparsity-driven noise. Nevertheless, the use of EM to include multimappers yielded only modest gains on young elements and never removed the fundamental ambiguity inherent to their repetitive sequences.

Our simulations used a deliberately stringent age threshold (2 million years) to define the “young” class, focusing on the most challenging loci. Such recent elements may represent only a minority of expressed TEs in some contexts. However, our results suggest that young age constitutes a systematic source of false positives in droplet-based scRNA-seq. To test hypothesis concerning young TEs, alternative experimental approaches (such as long-read scRNA-seq) should be considered. Alternatively, subfamily level analysis of young TE expression provides more robust quantification (at the cost of precluding analysis of the genomic context of individual insertions). The subfamily-level analyses presented here, however, should not be interpreted as a comprehensive benchmark of method performance, as we evaluated aggregation only after locus-level read assignment. Tools specifically designed for subfamily-level quantification in scRNA-seq, such as IRescue [24] and scTE [22], were not included because they do not provide locus-resolved counts. In addition, although SoloTE outputs subfamily-and family-level matrices, they only include reads not assignable to specific loci, which were excluded from our analysis. Consequently, its performance at these aggregation levels may differ from what is reported here.

Family-level analyses suggest substantial heterogeneity in quantification accuracy. Longer and more homogeneous families such as L1 and ERVL are particularly prone to misassignment, whereas SINEs (Alu, B2) are easier to resolve at the locus level. The difficulty in tracking expression of LINE insertions was already previously reported in both mouse [19] and human [20]. We hypothesize that in the case of LTR elements, the quantification could further be complicated by reads falling in the highly conserved long terminal repeat regions. These differences motivate family-aware quality control and tool selection strategies, depending on the families expected to be active in a given biological system. We note, however, that the simulations presented here do not capture all characteristics of real scRNA-seq data, most notably protocol-specific biases such as polyadenylation effects that may influence the representation of certain TE classes. Consequently, relative performance across TE families may differ in empirical datasets. The results should therefore be interpreted primarily as an indication of the challenges associated with resolving individual loci within subfamilies, assuming that the corresponding reads are detected.

Simulations containing both genes and TEs further revealed that gene-TE disambiguation is a major unresolved challenge. Tools vary in how they prioritize gene versus TE annotations during read assignment, resulting in false positives in both directions: gene-derived reads incorrectly assigned to TEs, and TE-derived reads incorrectly assigned to genes. Stellarscope is particularly sensitive to this issue because it retains reads overlapping gene bodies. However, all tools exhibited some degree of cross-assignment. While excluding gene-overlapping TEs a priori reduces false positives, it risks discarding genuinely expressed insertions, which is a nontrivial biological class. Improved joint gene-TE models and more refined annotation strategies will be needed to address these limitations.

We based our evaluation on a simulated ground truth, as no experimental method currently enables perfectly accurate quantification of transposable elements at the single cell level. While Splatter, like other scRNA-seq data simulation tools, may not fully capture the complexity of real single-cell count matrices and may introduce systematic biases [46], advancing simulation realism is itself still an active research area and beyond the scope of the present study. Here, our goal was instead to assess TE quantification accuracy under controlled, predefined expression levels. An important consequence of this design choice is that differences in relative abundance across loci, which may affect quantification performance, are not recapitulated. In future work, developing TE-aware simulation frameworks that better reflect the heterogeneity, abundance distributions and sequence characteristics of expressed TE loci would offer an interesting opportunity to further strengthen benchmarking efforts.

Beyond the tools evaluated here, it is important to note that locus-specific TE quantification remains a rapidly evolving area, with new methods frequently introduced and many existing tools still under active development. For example, MATES [25] is a recently proposed deep-learningbased approach that simultaneously models read-locus relationships and multi-mapping patterns to infer locus-level TE expression. Although promising, we were not able to include MATES in our benchmark because, at the time of analysis, open issues prevented its successful execution within our framework. As the method continues to mature and these software limitations are resolved, it will be straightforward to incorporate into our workflow. Indeed, our Snakemake-based bench-marking framework is modular and easy to extend, enabling future versions of MATES and any forthcoming tools to be added with minimal effort.

Here, we focused on short-read scRNA-seq because it remains the dominant technology and because existing TE tools have been designed around these data. However, short-read protocols impose fundamental limits on TE quantification due to their short fragment lengths. Long-read scRNA-seq has the potential to overcome these barriers by spanning insertion-specific mutations that distinguish copies within a family. Yet long-read assays still exhibit higher per-base error rates [47], which may counteract this advantage by introducing mapping uncertainty. Recent developments such as CELLO-seq [16] and error-corrected long-read protocols [48] provide promising directions, although they have so far been applied only to a limited number of datasets. Integrating these emerging experimental innovations with dedicated computational approaches will be essential for moving beyond the limitations of current short-read workflows.

Overall, our findings define the practical boundaries of current locus-specific TE quantification and emphasize the need for improved computational and experimental methods, particularly for young, highly repetitive elements and for accurate gene-TE disambiguation.

## 4 Conclusions

Our benchmark provides a unified, ground-truth–validated assessment of current tools for locus-specific TE quantification in short-read scRNA-seq. Three conclusions are robust across datasets and simulation settings. First, locus-level quantification is reliable for evolutionarily older elements, but young, highly repetitive TEs remain intrinsically difficult to resolve with short reads. Second, multi-mapper handling offers limited improvements: methods based on EM improve precision modestly but reduce sensitivity, and unique-mapper strategies often perform comparably while producing fewer false positives. Third, gene-TE misassignment is common and must be explicitly checked, as errors occur in both directions and can strongly confound biological interpretation.

These findings motivate several best-practice recommendations, which we include below:

(i) Use locus-level quantification for older elements, while the interpretation of results from young-locus calls needs caution. (ii) Prefer unique-mapper strategies (e.g., SoloTE default) when maximizing precision for locus calls is essential. If including multi-mappers, apply EM (e.g., Stellarscope) with conservative posterior filters and report sensitivity loss. (iii) For discovery or atlas-scale surveys where TE identity at base resolution is less critical, aggregate counts at the subfamily level to improve robustness (especially for young elements). (iv) Incorporate explicit gene-TE overlap checks into the analysis pipeline and report the proportion of TE calls overlapping expressed genes. Our Snakemake workflow and all simulated datasets are publicly available for full reproducibility and extension. As long-read scRNA-seq and new TE-aware protocols mature, integrating them with improved computational models will be critical for unlocking truly locus-resolved TE biology in single cells. This benchmark establishes a foundation for such future developments and provides a reference framework for evaluating forthcoming tools and sequencing technologies.

## 5 Methods

### 5.1 Datasets

The mESC dataset was obtained from Iturbide et al. [34]. We selected the sample that was exposed to retinoic acid for 48h, which has the largest population of 2CLCs. The mouse olfactory mucosa dataset was published by Horgue et al. [37]. The PBMC dataset was downloaded from the 10XGenomics database [40] and contains around 8 thousand cells from a healthy donor’s blood sample. All the datasets were sequenced with the Chromium Single Cell 3’ v3 10X Genomics protocol. Details in Table S2.

### 5.2 TE age estimation

To estimate TE age, we used milli-divergence values (*p_AB_*) from RepeatMasker annotations and applied the Jukes–Cantor model to compute evolutionary distances (*d_AB_* = —(3*/*4) ln (1 — (4*/*3)*p_AB_*). Following Berrens et al., 2022 [16], distances were then converted to millions of years using a per-site, per-year substitution rate (*µ*) of 2.2 × 10*^—^*^9^ for human and 4.5 × 10*^—^*^9^ for mouse: age (mya) = (*d_AB_* * 100)*/*(2*µ* * 100) × 10^3^.

### 5.3 Synthetic data generation

The Minnow framework (v0.1.0, [43]) was used to generate simulated scRNA-seq datasets (Fig. 2), mimicking gene and TE-level multi-mappability of the mESC datasets described above whilst considering realistic PCR amplification bias and sequencing errors. We first generated TE nucleotide sequence files from regions annotated in the TE GTF files curated and published by the Hammell lab (mm10 version, [49]) and used these as custom transcriptomes to build Salmon (v1.10.3, [50]) indexes. We then ran Alevin [51] (integrated with the Salmon software) on the 10X mESC dataset to produce the BFH files required by Minnow. Indexing and estimation were performed using default parameters.

scRNA-seq count matrices containing 500 cells each were simulated using Splatter (v1.18.2, [42]), with the Splat model (based on a gamma-Poisson distribution), with default parameters. We selected loci that passed Minnow’s filters during the indexing step, and subsampled them to ensure a balanced representation of TE families. Specifically, we selected up to 100 old and 300 young TEs per family, retaining only families with at least 50 loci. This resulted in a total of 3,173 old and 2,100 young loci. 2,000 highly variable genes were also selected (see Section 5.6). The selected features were then randomly assigned to the rows of the simulated matrices (all configurations shown in Fig. 2B).

The Minnow simulation step was subsequently run on the Splatter matrices to generate matching 101 bp-long sequencing reads. In downstream analyses, the simulated datasets were treated as unstranded, since we found that approximately half of the features had reads mapping to the same strand as the originating feature, while the remainder mapped to the opposite strand.

### 5.4 scRNA-seq preprocessing

The scRNA-seq analysis pipeline, including pre-processing and gene and TE quantification, was implemented as a modular Snakemake (v8.18.2, [29]) workflow, designed to be easily expanded to new datasets and tools. Our pipeline is publicly available on GitHub (https://github.com/ScialdoneLab/TEbenchmarking). To ensure reproducibility, separate Conda (v24.7.1, [52]) environments were used for each Snakemake rule, and a Docker (v24.0.2, [53]) container was also generated for the whole workflow.

Alignment of the scRNA-seq datasets, both real and synthetic, was performed using STARsolo [31] (implemented within STAR, v2.7.11, [54]). GENCODE genome assemblies (GRCh38.p14 for human and GRCm38.p4 for mouse) and gene annotations (release 30 for human and M11 for mouse) were used for indexing. STARsolo was then run with multimapping options--winAnchorMultimapNmax 100 and --outFilterMultimapNmax 100, and filtering was applied using a minimum mapping quality score of 30.

### 5.5 TE quantification

SoloTE (v1.09, [18]) was run with default parameters on alignment files in which only the best alignment for each read was retained (using STAR’s –outSAMmultNmax 1 option). To assess the impact of including multimapping reads, we modified the SoloTE source code to expose the internal MAPQ threshold (used to distinguish locus-level from family-level quantification) as a tunable parameter. We then replaced the default MAPQ cutoff of 255 (unique alignments only) with progressively lower thresholds (2, 1, and 0), thereby allowing inclusion of reads mapping to at most 2 loci, at most 4 loci, or to any number of loci, respectively (Table S1). SoloTE was executed separately under each configuration.

Stellarscope (v1.4, [23]) was run using the TE GTF files provided by the Hammell lab, which derive from RepeatMasker [55] annotations (version mm10 for mouse and hg38 for human datasets [49]). The “pseudobulk” pooling mode and the “best exclude” reassignment mode were used. For real datasets, we also tested the “celltype” pooling mode by providing manually curated cell-type annotations based on marker gene expression. For simulated data, we additionally tested the “conf” reassignment mode, including only UMIs assigned with a posterior probability score higher than 0.9.

To quantify transposons directly with STARsolo, we replaced the GTF files for gene annotation with the TE annotation GTF files [49]. Mapping was performed with and without the EM algorithm enabled via the –soloMultiMappers EM option.

### 5.6 Downstream analysis of gene and TE count matrices

The Seurat R package (v5.1.0, [56]) was used for downstream analysis of scRNA-seq count matrices. The gene and TE count matrices were analysed separately.

Only cells passing STARsolo’s quality filters were kept for downstream analysis. Features expressed in less than 2% of cells and loci classified as “Other”, “Satellite”, “Unknown” or “RNA” in the RepeatMasker annotation (less than 0.1% of loci in all samples) were removed from all count matrices. Feature counts (of both TEs and genes) were normalized by dividing them by the total counts for that cell and multiplying them by a scaling factor 10,000, and then natural-log transformed. Highly variable features were selected with the “vst” method implemented in Seurat. The top 4,000 highly variable genes and 4,000 highly variable TEs were selected for downstream tasks in all datasets. The top 2,000 highly variable genes from the mESC dataset were used for mixed gene and TE simulation (Fig. 2B, see Section 5.3).

Principal Component Analysis (PCA) was computed using only highly variable genes. KNN graph was then computed using local neighborhood of 20 cells. Cells were then clustered using the Leiden algorithm. To identify comparable clustering resolutions for gene- and TE-derived analyses, we performed clustering across a range of resolution parameters from 0.5 to 2.0 in increments of 0.1. For each resolution, we generated cluster assignments and computed the ARI between all pairs of resulting clusterings. The resolutions that maximized the ARI were selected for downstream comparisons. For visualisation, UMAP (Uniform Manifold Approximation and Projection)[57] representations were built with default parameters.

Cell type annotation was manually performed for all datasets based on the expression of well-characterized marker genes (Tables S3, S4, S5). The human PBMC cluster was then randomly subsampled within each cell type to ensure the same cell number across all defined types. This approach prevented more abundant cell types from biasing the results of the EM algorithms implemented in Stellarscope and STARsolo, and allowed for an unbiased evaluation of cell type label purity within the TE-derived clusters. One PBMC cluster of 229 cells could not be reliably annotated and was therefore excluded from further analysis. All gene and TE quantification tools were then rerun on the subsampled dataset.

Purity of the cell type labels shown in Fig. 1 was computed for each cell as the fraction of cells of the same type among its 20 nearest neighbors in the KNN graph.

### 5.7 Evaluation metrics and statistical tests

For each cell, we computed precision as *TP/*(*TP* + *FP*) detected loci, recall as *TP/*(*TP* + *FN*) detected loci, F1 score as 2(*Precision* * *Recall*)*/*(*Precision* + *Recall*). Metrics were then averaged across cells.

Spearman’s correlation coefficients were computed between simulated and estimated counts, after normalization (using the same approach as in Section 5.6).

To identify locus-level features associated with higher correlation accuracy, we fit a generalized linear mixed-effects model with a binomial distribution and logit link (using the lme4 R package, v.1.1-37 [58]). For each locus, the binary outcome *y_ij_* indicated whether the correlation exceeded the median across loci (high accuracy = 1, low accuracy = 0). Standardized TE length (*x*_1*,ij*_) and average simulated expression (*x*_2*,ij*_) were included as fixed effects, and TE quantification tool was included as a random intercept *b_j_*:

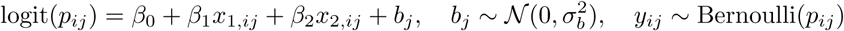

## 6 Declarations

### Availability of data and materials

The mouse ESC data [34] is available under ArrayExpress accession no. E-MTAB-8869. The mouse olfactory mucosa scRNA-seq data [37] is deposited in the NCBI GEO database under accession code GSE185168. The human PBMC dataset can be downloaded from the 10X Genomics website [40].

The simulated datasets generated for this benchmarking are available on Zenodo (10.5281/zenodo.18772673). The snakemake workflow and notebooks used for tool evaluation and data simulation are available on Github (https://github.com/ScialdoneLab/TEbenchmarking).

## Acknowledgments

We thank Maria Elena Torres Padilla, Ines Hellmann, Johanna Klughammer, Timothy Ramnarine, Simon Mages, and all members of the Scialdone and Vallejos labs for feedback and insightful discussions. This work was supported by the Helmholtz Association (AS, VF). VF is supported by the International Helmholtz-Edinburgh Research School for Epigenetics (EpiCrossBorders). This work was supported by the BMBF-funded de.NBI Cloud within the German Network for Bioinformatics Infrastructure (de.NBI), to which A.S. and V.F. were granted access.

## Authors’ contributions

A.S. and C.V. conceived and jointly supervised the project. V.F. performed all the computational analyses. Manuscript writing – Original Draft: V.F.; Review & Editing: V.F., C.V., A.S.; Funding Acquisition: A.S. and C.V.. All authors read and approved the final manuscript.

## References

[1] Jonathan N Wells and Cédric Feschotte. “A field guide to eukaryotic transposable elements.” In: Annu Rev Genet 54 (Nov. 2020), pp. 539–561. doi: 10.1146/annurev-genet-040620-022145. (visited on 11/02/2023).

[2] Diego Rodriguez-Terrones and Maria-Elena Torres-Padilla. “Nimble and ready to mingle: transposon outbursts of early development.” In: Trends Genet 34.10 (Oct. 2018), pp. 806– 820. issn: 01689525. doi: 10.1016/j.tig.2018.06.006. url: https://linkinghub.elsevier.com/retrieve/pii/S0168952518301197 (visited on 11/02/2023).

[3] Camille Ravel-Godreuil, Rania Znaidi, Tom Bonnifet, Rajiv L Joshi, and Julia Fuchs. “Transposable elements as new players in neurodegenerative diseases”. en. In: FEBS Lett 595.22 (Oct. 2021), pp. 2733–2755.

[4] Kathleen H“”Burns. “”Transposable elements in cancer””. en. In: Nat Rev Cancer 17.7 (June 2017), pp. 415–424.

[5] Guillaume Bourque et al. “Ten things you should know about transposable elements.” In: Genome Biol 19.1 (Nov. 2018), p. 199. issn: 1474-760X. doi: 10.1186/s13059-018-1577-z. url: https://genomebiology.biomedcentral.com/articles/10.1186/s13059-018-1577-z (visited on 05/21/2024).

[6] Diwash Jangam, Cédric Feschotte, and Esther Betŕan. “Transposable element domestication as an adaptation to evolutionary conflicts.” In: Trends Genet 33.11 (Nov. 2017), pp. 817–831. doi: 10.1016/j.tig.2017.07.011. (visited on 12/23/2025).

[7] Yann Bourgeois and Stéphane Boissinot. “On the population dynamics of junk: A review on the population genomics of transposable elements.” In: Genes 10.6 (May 2019). doi: 10.3390/genes10060419. (visited on 01/16/2026).

[8] Vasavi Sundaram and Joanna Wysocka. “Transposable elements as a potent source of diverse cis-regulatory sequences in mammalian genomes”. In: Philosophical Transactions of the Royal Society B: Biological Sciences 375.1795 (Feb. 2020), p. 20190347. issn: 0962-8436. doi: 10.1098/rstb.2019.0347. eprint: https://royalsocietypublishing.org/rstb/article-pdf/doi/10.1098/rstb.2019.0347/258565/rstb.2019.0347.pdf.

[9] Luisa Di Stefano. “All Quiet on the TE Front? The Role of Chromatin in Transposable Element Silencing”. In: Cells 11.16 (2022). issn: 2073-4409. doi: 10.3390/cells11162501. url: https://www.mdpi.com/2073-4409/11/16/2501.

[10] Gireesh K. Bogu, Ferran Reverter, Marc A. Marti-Renom, Michael P. Snyder, and Roderic Guigo’. “Atlas of transcriptionally active transposable elements in human adult tissues”. In: bioRxiv (2019). doi: 10.1101/714212. eprint: https://www.biorxiv.org/content/early/2019/07/24/714212.full.pdf. url: https://www.biorxiv.org/content/early/2019/07/24/714212.

[11] İbrahim Avşar Ilık, Xu Yang, Z Z Zhao Zhang, and Tuğçe Aktaş. “Transcriptional and post-transcriptional regulation of transposable elements and their roles in development and disease.” In: Nat Rev Mol Cell Biol 26.10 (Oct. 2025), pp. 759–775. doi: 10.1038/s41580-025-00867-8. (visited on 01/02/2026).

[12] Edward B Chuong, Nels C Elde, and Cédric Feschotte. “Regulatory activities of transposable elements: from conflicts to benefits.” In: Nat Rev Genet 18.2 (Feb. 2017), pp. 71–86. doi: 10.1038/nrg.2016.139. (visited on 11/21/2016).

[13] Martin Proks, Nazmus Salehin, and Joshua M. Brickman. “Deep learning-based models for preimplantation mouse and human embryos based on single-cell RNA sequencing”. In: Nat Methods (Nov. 2024). issn: 1548-7091. doi: 10.1038/s41592-024-02511-3. url: https://www.nature.com/articles/s41592-024-02511-3 (visited on 11/28/2024).

[14] Cheng Zhao et al. “A comprehensive human embryo reference tool using single-cell RNA-sequencing data.” In: Nat Methods 22.1 (Jan. 2025), pp. 193–206. issn: 1548-7091. doi: 10.1038/s41592-024-02493-2. url: https://www.nature.com/articles/s41592-024-02493-2 (visited on 11/25/2024).

[15] Sophie Lanciano and Gael Cristofari. “Measuring and interpreting transposable element expression.” In: Nat Rev Genet 21.12 (Dec. 2020), pp. 721–736. issn: 1471-0056. doi: 10.1038/s41576-020-0251-y. url: http://www.nature.com/articles/s41576-020-0251-y (visited on 06/29/2020).

[16] Rebecca V Berrens et al. “Locus-specific expression of transposable elements in single cells with CELLO-seq.” In: Nat Biotechnol 40.4 (Apr. 2022), pp. 546–554. issn: 1087-0156. doi: 10.1038/s41587-021-01093-1. url: https://www.nature.com/articles/s41587-021-01093-1 (visited on 11/15/2021).

[17] Emmanuelle Lerat. “Recent bioinformatic progress to identify epigenetic changes associated to transposable elements.” In: Front Genet 13 (May 2022), p. 891194. doi: 10.3389/fgene.2022.891194. (visited on 01/18/2026).

[18] Rocío Rodríguez-Quiroz and Braulio Valdebenito-Maturana. “SoloTE for improved analysis of transposable elements in single-cell RNA-Seq data using locus-specific expression.” In: Commun Biol 5.1 (Oct. 2022), p. 1063. doi: 10.1038/s42003-022-04020-5. (visited on 11/08/2023).

[19] Robert Schwarz, Philipp Koch, Jeanne Wilbrandt, and Steve Hoffmann. “Locus-specific expression analysis of transposable elements.” In: Brief Bioinformatics 23.1 (Jan. 2022). doi: 10.1093/bib/bbab417. (visited on 10/30/2023).

[20] Natalia Savytska, Peter Heutink, and Vikas Bansal. “Transcription start site signal profiling improves transposable element RNA expression analysis at locus-level.” In: Front Genet 13 (Oct. 2022), p. 1026847. doi: 10.3389/fgene.2022.1026847. (visited on 07/10/2024).

[21] Kathryn O’Neill, David Brocks, and Molly Gale Hammell. “Mobile genomics: tools and techniques for tackling transposons”. In: Philosophical Transactions of the Royal Society B: Biological Sciences 375.1795 (Feb. 2020), p. 20190345. issn: 0962-8436. doi: 10.1098/rstb.2019.0345. eprint: https://royalsocietypublishing.org/rstb/article-pdf/doi/10.1098/rstb.2019.0345/258803/rstb.2019.0345.pdf.

[22] Jiangping He et al. “Identifying transposable element expression dynamics and heterogeneity during development at the single-cell level with a processing pipeline scTE.” In: Nat Commun 12.1 (Mar. 2021), p. 1456. doi: 10.1038/s41467-021-21808-x. (visited on 11/08/2023).

[23] Helena Reyes-Gopar et al. “A single-cell transposable element atlas of human cell identity”. In: Cell Reports Methods 5.7 (July 2025).

[24] Benedetto Polimeni, Federica Marasca, Valeria Ranzani, and Beatrice Bodega. “IRescue: uncertainty-aware quantification of transposable elements expression at single cell level.” In: Nucleic Acids Res 52.19 (Oct. 2024), e93. doi: 10.1093/nar/gkae793.(visited on 01/13/2025).

[25] Ruohan Wang, Yumin Zheng, Zijian Zhang, Kailu Song, Erxi Wu, Xiaopeng Zhu, Tao P Wu, and Jun Ding. “MATES: a deep learning-based model for locus-specific quantification of transposable elements in single cell.” In: Nat Commun 15.1 (Oct. 2024), p. 8798. doi: 10.1038/s41467-024-53114-7. (visited on 01/22/2025).

[26] Anthony Sonrel et al. “Meta-analysis of (single-cell method) benchmarks reveals the need for extensibility and interoperability.” In: Genome Biol 24.1 (May 2023), p. 119. issn: 1474-760X. doi: 10.1186/s13059-023-02962-5. url: https://genomebiology.biomedcentral.com/ articles/10.1186/s13059-023-02962-5 (visited on 03/07/2024).

[27] Jorge A. Tzec-Interián, Daianna González-Padilla, and Elsa B. Góngora-Castillo. “Bioinformatics perspectives on transcriptomics: A comprehensive review of bulk and single-cell RNA sequencing analyses”. In: Quant Biol 13.2 (June 2025). issn: 2095-4689. doi: 10.1002/qub2.78. url: https://onlinelibrary.wiley.com/doi/10.1002/qub2.78 (visited on 02/05/2026).

[28] Valentine Svensson, Eduardo da Veiga Beltrame, and Lior Pachter. “A curated database reveals trends in single-cell transcriptomics”. In: Database 2020 (Nov. 2020), baaa073. issn: 1758-0463. doi:10.1093/database/baaa073. eprint: https://academic.oup.com/database/article-pdf/doi/10.1093/database/baaa073/34568703/baaa073.pdf.

[29] Felix Mölder, et al. “Sustainable data analysis with Snakemake”. In: F1000Res 10 (Jan. 2021), p. 33. issn: 2046-1402. doi: 10.12688/f1000research.29032.1. url: https://f1000research.com/articles/10-33/v1 (visited on 01/17/2022).

[30] Matthew L Bendall et al. “Telescope: Characterization of the retrotranscriptome by accurate estimation of transposable element expression.” In: PLoS Comput Biol 15.9 (Sept. 2019), e1006453. doi: 10.1371/journal.pcbi.1006453. (visited on 08/18/2020).

[31] Benjamin Kaminow, Dinar Yunusov, and Alexander Dobin. “STARsolo: accurate, fast and versatile mapping/quantification of single-cell and single-nucleus RNA-seq data”. In: bioRxiv (2021). doi: 10.1101/2021.05.05.442755. eprint: https://www.biorxiv.org/content/early/2021/05/05/2021.05.05.442755.full.pdf. url: https://www.biorxiv.org/content/early/2021/05/05/2021.05.05.442755.

[32] Aaron R Quinlan and Ira M Hall. “BEDTools: a flexible suite of utilities for comparing genomic features.” In: Bioinformatics 26.6 (Mar. 2010), pp. 841–842. doi: 10.1093/bioinformatics/btq033. (visited on 11/18/2016).

[33] Konstantin Tretyakov. intervaltree. Python package. url: https://pypi.org/project/intervaltree/.

[34] Ane Iturbide, Mayra L Ruiz Tejada Segura, Camille Noll, Kenji Schorpp, Ina Rothenaigner, Elias R Ruiz-Morales, Gabriele Lubatti, Ahmed Agami, Kamyar Hadian, Antonio Scialdone, and Maria-Elena Torres-Padilla. “Retinoic acid signaling is critical during the totipotency window in early mammalian development.” In: Nat Struct Mol Biol 28.6 (June 2021), pp. 521–532. issn: 1545-9993. doi: 10.1038/s41594-021-00590-w. url: http://www.nature.com/articles/s41594-021-00590-w (visited on 02/09/2025).

[35] Youjia Guo, Ten D Li, Andrew J Modzelewski, and Haruhiko Siomi. “Retrotransposon renaissance in early embryos.” In: Trends Genet 40.1 (Jan. 2024), pp. 39–51. issn: 01689525. doi: 10.1016/j.tig.2023.10.010. url: https://linkinghub.elsevier.com/retrieve/ pii/%7BS016895252300238X%7D (visited on 07/07/2024).

[36] Lauryn A Deaville and Rebecca V Berrens. “Technology to the rescue: how to uncover the role of transposable elements in preimplantation development.” In: Biochem Soc Trans 52.3 (June 2024), pp. 1349–1362. doi: 10.1042/{BST20231262}. url: 10.1042/%7BBST20231262%7D (visited on 05/22/2024).

[37] Luis Flores Horgue, Alexis Assens, Leon Fodoulian, Leonardo Marconi, Jöel Tuberosa, Alexander Haider, Madlaina Boillat, Alan Carleton, and Ivan Rodriguez. “Transcriptional adaptation of olfactory sensory neurons to GPCR identity and activity.” In: Nature Communications 13.1 (May 2022), p. 2929. issn: 2041-1723. doi: 10.1038/s41467-022-30511-4. url: https://www.nature.com/articles/s41467-022-30511-4 (visited on 06/17/2024).

[38] Xinmin Zhang and Stuart Firestein. “The olfactory receptor gene superfamily of the mouse.” In: Nature Neuroscience 5.2 (Feb. 2002), pp. 124–133. issn: 1097-6256. doi: 10.1038/nn800. (visited on 05/20/2021).

[39] Ariel D Pourmorady et al. “RNA-mediated symmetry breaking enables singular olfactory receptor choice.” In: Nature 625.7993 (Jan. 2024), pp. 181–188. issn: 0028-0836. doi: 10.1038/s41586-023-06845-4. url: https://www.nature.com/articles/s41586-023-06845-4 (visited on 12/20/2023).

[40] *8k PBMCs from a Healthy Donor, Universal 3’ dataset*. url: https://www.10xgenomics.com/datasets/8-k-pbm-cs-from-a-healthy-donor-2-standard-2-1-0.

[41] Malte D Luecken et al. “Defining and benchmarking open problems in single-cell analysis.” In: Nature Biotechnology 43.7 (July 2025), pp. 1035–1040. issn: 1087-0156. doi: 10.1038/s41587-025-02694-w. url: https://www.nature.com/articles/s41587-025-02694-w (visited on 01/23/2026).

[42] Luke Zappia, Belinda Phipson, and Alicia Oshlack. “Splatter: simulation of single-cell RNA sequencing data.” In: Genome Biol 18.1 (Sept. 2017), p. 174. doi: 10.1186/s13059-017-1305-0. (visited on 07/25/2017).

[43] Hirak Sarkar, Avi Srivastava, and Rob Patro. “Minnow: a principled framework for rapid simulation of dscRNA-seq data at the read level”. In: Bioinformatics 35.14 (July 2019), pp. i136– i144. issn: 1367-4803. doi: 10.1093/bioinformatics/btz351. eprint: https://academic.oup.com/bioinformatics/articlepdf/35/14/i136/50721031/bioinformatics\_35\_14\_i136.pdf.

[44] W. N. Venables and B. D. Ripley. Modern Applied Statistics with S. Fourth. ISBN 0-387-95457-0. New York: Springer, 2002. url: https://www.stats.ox.ac.uk/pub/MASS4/.

[45] H. Läuter. “Silverman, B. W.: Density Estimation for Statistics and Data Analysis. Chapman & Hall, London – New York 1986, 175 pp., £12.—”. In: Biometrical Journal 30.7 (1988), pp. 876–877. doi: 10.1002/bimj.4710300745. eprint: https://onlinelibrary.wiley.com/doi/pdf/10.1002/bimj.4710300745. url: https://onlinelibrary.wiley.com/doi/abs/10.1002/bimj.4710300745.

[46] Helena L Crowell, Sarah X Morillo Leonardo, Charlotte Soneson, and Mark D Robinson. “The shaky foundations of simulating single-cell RNA sequencing data.” In: Genome Biol 24.1 (Mar. 2023), p. 62. doi: 10.1186/s13059-023-02904-1. (visited on 02/14/2024).

[47] Rebekah K Loving et al. “Long-read sequencing transcriptome quantification with lr-kallisto.” In: PLoS Comput Biol 21.12 (Dec. 2025), e1013692. doi: 10.1371/journal.pcbi.1013692. (visited on 01/30/2026).

[48] Xuemei Li, Keying Lu, Xiao Chen, Kailing Tu, and Dan Xie. “capTEs enables locus-specific dissection of transcriptional outputs from reference and nonreference transposable elements.” In: Commun Biol 6.1 (Sept. 2023), p. 974. doi: 10.1038/s42003-023-05349-1. (visited on 02/28/2024).

[49] HammellLab. *Curated TE annotation GTF files*. url: https://www.dropbox.com/scl/fo/jdpgn6fl8ngd3th3zebap/ACdZkShDC1au-OckIipI5kM/TEtranscripts/TE_GTF?dl=0&rlkey=41oz6ppggy82uha5i3yo1rnlx&subfolder_nav_tracking=1.

[50] Rob Patro, Geet Duggal, Michael I Love, Rafael A Irizarry, and Carl Kingsford. “Salmon provides fast and bias-aware quantification of transcript expression.” In: Nat Methods 14.4 (Apr. 2017), pp. 417–419. doi: 10.1038/nmeth.4197. (visited on 06/24/2019).

[51] Avi Srivastava, Laraib Malik, Tom Smith, Ian Sudbery, and Rob Patro. “Alevin efficiently estimates accurate gene abundances from dscRNA-seq data.” In: Genome Biol 20.1 (Mar. 2019), p. 65. issn: 1474-760X. doi: 10.1186/s13059-019-1670-y. url: https://genomebiology.biomedcentral.com/articles/10.1186/s13059-019-1670-y (visited on 04/28/2019).

[52] *Anaconda Software Distribution. Computer software.* Version 2-2.4.0. 2016. url: https://anaconda.com.

[53] Dirk Merkel. “Docker: lightweight linux containers for consistent development and deployment”. In: Linux journal 2014.239 (2014), p. 2.

[54] Alexander Dobin, Carrie A Davis, Felix Schlesinger, Jorg Drenkow, Chris Zaleski, Sonali Jha, Philippe Batut, Mark Chaisson, and Thomas R Gingeras. “STAR: ultrafast universal RNA-seq aligner.” In: Bioinformatics 29.1 (Jan. 2013), pp. 15–21. doi:10.1093/bioinformatics/bts635. (visited on 04/25/2016).

[55] Maja Tarailo-Graovac and Nansheng Chen. “Using RepeatMasker to identify repetitive elements in genomic sequences.” In: Curr Protoc Bioinformatics Chapter 4 (Mar. 2009), Unit 4.10. doi: 10.1002/0471250953.bi0410s25. (visited on 10/26/2023).

[56] Yuhan Hao, Tim Stuart, Madeline H Kowalski, Saket Choudhary, Paul Hoffman, Austin Hartman, Avi Srivastava, Gesmira Molla, Shaista Madad, Carlos Fernandez-Granda, and Rahul Satija. “Dictionary learning for integrative, multimodal and scalable single-cell analysis”. In: Nature Biotechnology (2023). doi: 10.1038/s41587-023-01767-y.

[57] Leland McInnes, John Healy, Nathaniel Saul, and Lukas Großberger. “UMAP: Uniform Manifold Approximation and Projection”. In: Journal of Open Source Software 3.29 (2018), p. 861. doi: 10.21105/joss.00861.

[58] Douglas Bates, Martin Mächler, Ben Bolker, and Steve Walker. “Fitting Linear Mixed-Effects Models Using lme4”. In: Journal of Statistical Software 67.1 (2015), pp. 1–48. doi: 10.18637/jss.v067.i01.

