## Supplementary Figures and Tables for "Benchmarking computational tools for locus-specific analysis of transposable elements in single-cell RNA-seq datasets"

### 1 Supplementary Figures and Tables

Table S1: Possible configurations of our modified SoloTE implementation to control the inclusion of multimapping reads in the benchmarking analysis. Each configuration corresponds to a different minimum MAPQ cutoff determining which reads are retained for locus-specific quantification.

| **Configuration** | **Read inclusion criteria** | **MAPQ threshold** |
|---|---|---|
| SoloTE unique | Only uniquely mapping reads included | 255 |
| SoloTE max2loci | Reads mapping to at least 2 loci included | 2 |
| SoloTE max4loci | Reads mapping to at least 4 loci included | 1 |
| SoloTE multi | All multimapping reads included | 0 |

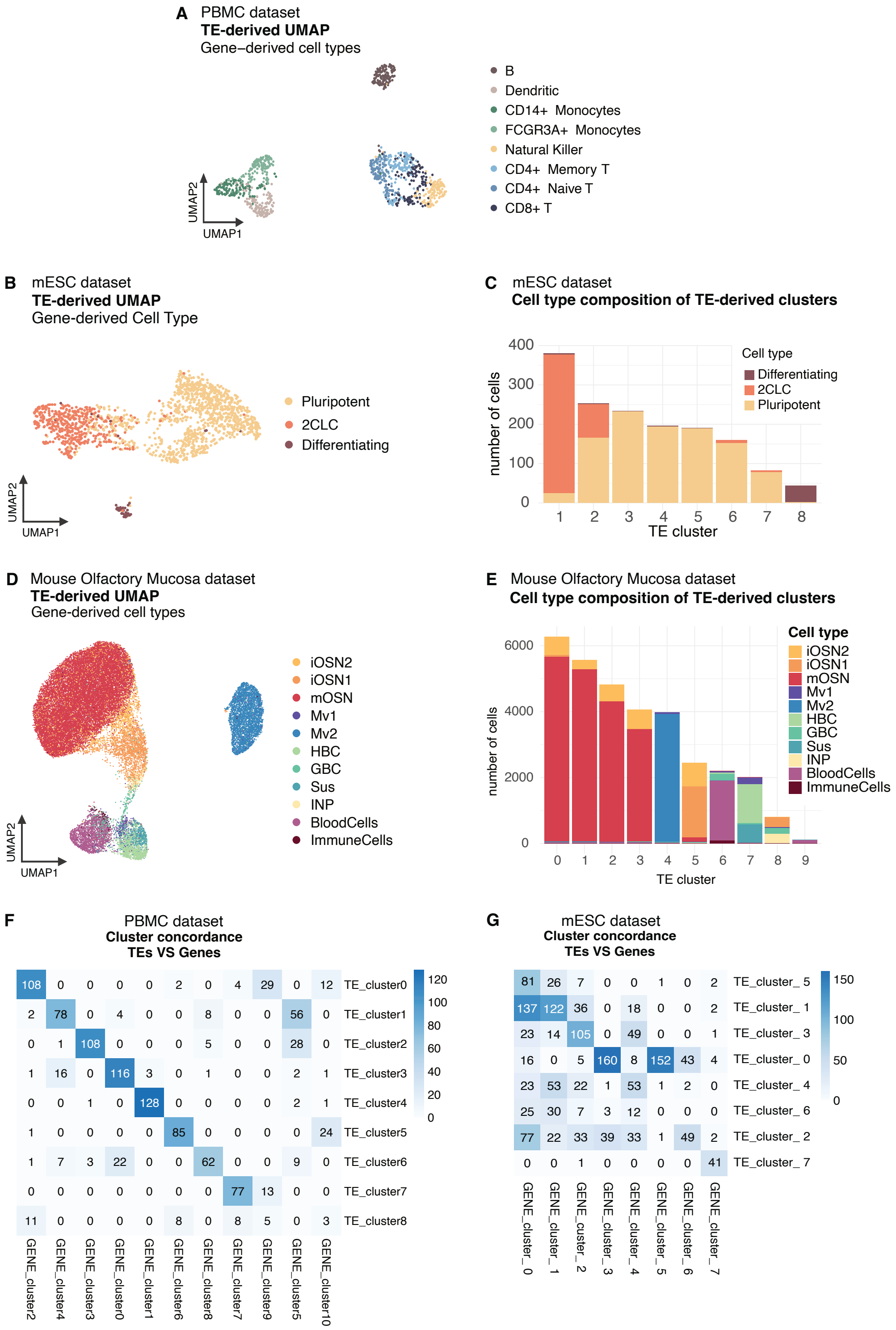


Figure S1: **A** UMAP of the human PBMC dataset, derived from SoloTE's locus TE count matrix, colored by gene-derived cell type annotation. **B** UMAP of the mESC dataset, derived from SoloTE's locus TE count matrix, colored by gene-derived cell type annotation. **C** Number of cells assigned to each gene-derived cell type within each TE-derived cluster in the mESC dataset. **D** UMAP of the mouse olfactory mucosa dataset, derived from SoloTE's locus TE count matrix, colored by gene-derived cell type annotation. **E** Number of cells assigned to each TE- (rows) and gene- (columns) derived Leiden cluster in the human PBMC dataset. Clusters were obtained with the resolutions that maximize the ARI between the two clusterings (ARI=0.624). **F** Number of cells assigned to each TE- (rows) and gene- (columns) derived Leiden cluster in the mESC dataset. Clusters were obtained with the resolutions that maximize the ARI between the two clusterings (ARI=0.230).

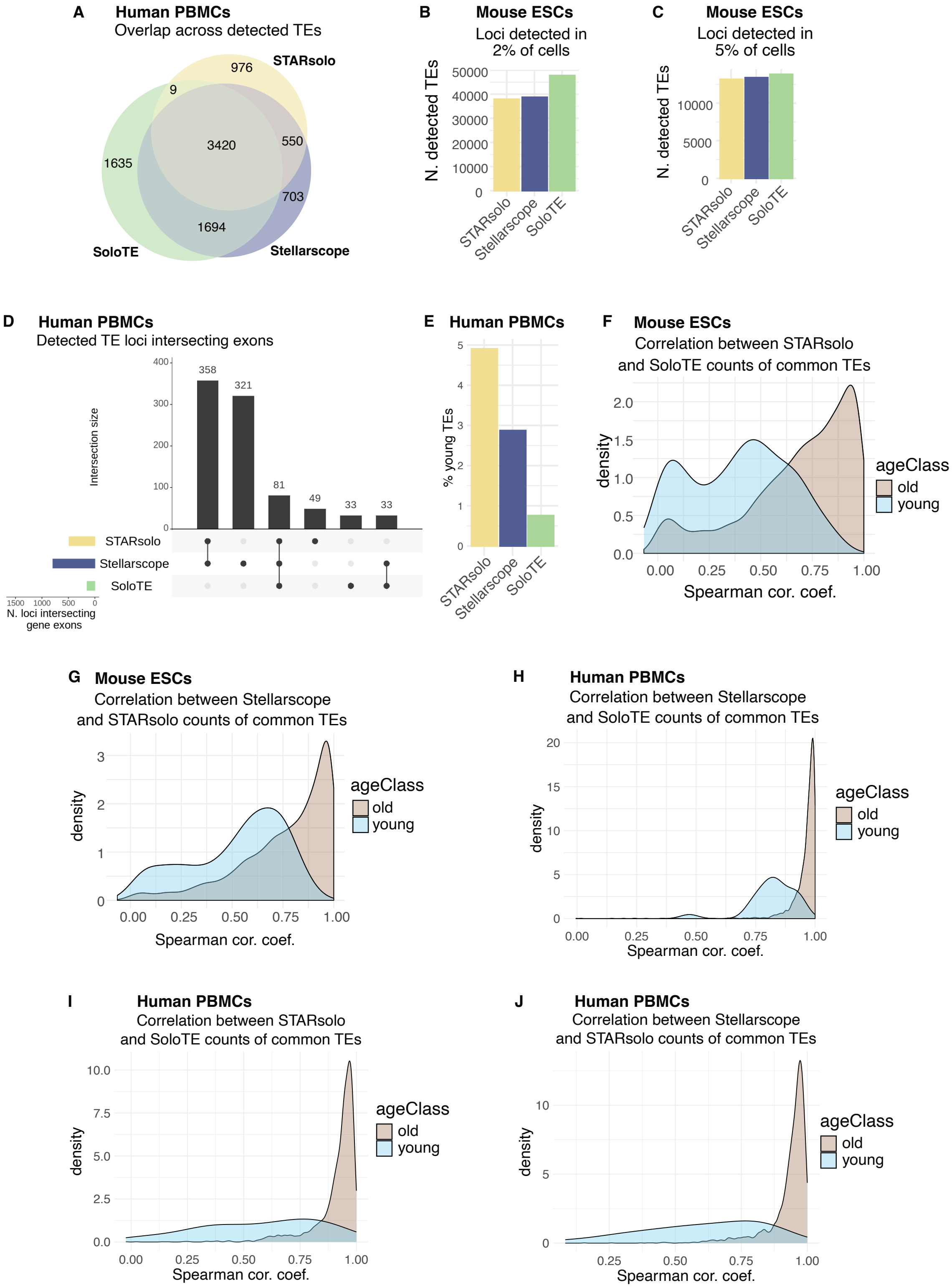


Figure S2: **A** Venn diagram showing the overlaps of detected TE loci across tools in the human PBMC dataset. **B** Total number of TE loci detected by each tool in at least 2% of cells in the mouse ESC dataset. **C** Total number of TE loci detected by each tool in at least 5% of cells in the mouse ESC dataset. **D** Upset plot of expressed TE loci intersecting gene exons identified by each tool in the human PBMC dataset. **E** Percentage of young TEs among loci detected in the PBMC dataset by each tool. **F,G** Distribution of Spearman correlation coefficients between normalized counts obtained with STARsolo and SoloTE (F), and between STARsolo and Stellarscope (G) in the mouse ESC dataset, calculated per locus across cells. Only loci detected by all evaluated tools are included. **H,I,J** Distribution of Spearman correlation coefficients between normalized counts obtained with SoloTE and Stellarscope (H), STARsolo and SoloTE (I), and STARsolo and Stellarscope (J) in the PBMC dataset, calculated per locus across cells. Only loci detected by all tools are included.

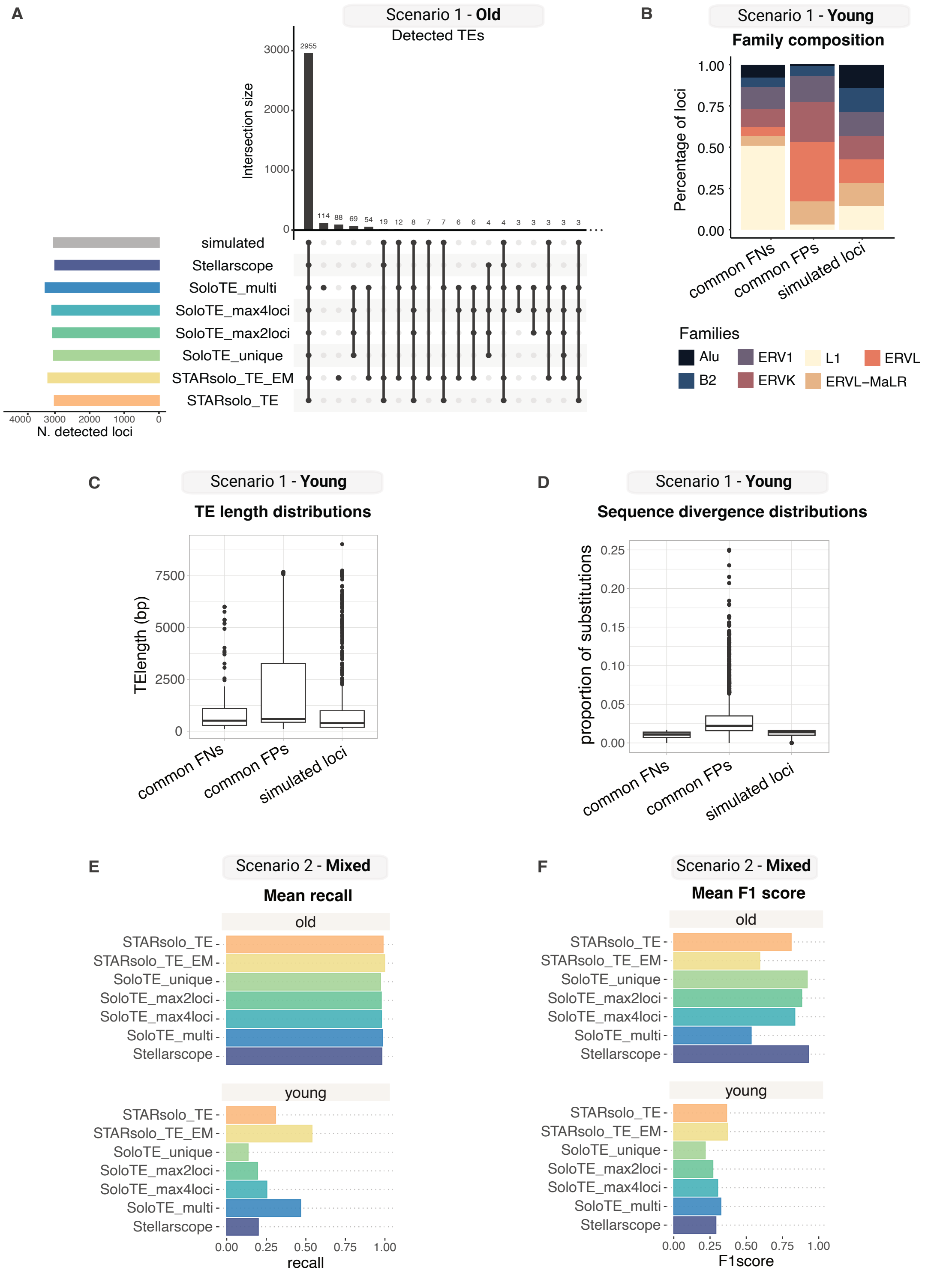


Figure S3: **A** Upset plot of TE loci present in the old TE simulated matrix (grey bar) and detected by each tool. SoloTE configurations are described in Sec. 5.5. **B** Percentage of loci belonging to each TE family in the sets of loci not detected by any tool but present in the simulation ("common FNs"), wrongly detected by all tools ("common FPs") and present in the simulated ground truth ("simulated loci"). **C** Distribution of age of common FNs, common FPs and simulated loci. **D** Distribution of proportion of substitutions from family consensus of common FNs, common FPs and simulated loci. **E,F** Recall (E) and F1 score (F) computed per cell and then averaged across cells, using the binarized count matrix obtained from the simulation containing both old and young loci (Scenario 2). Results are shown separately for old and young loci.

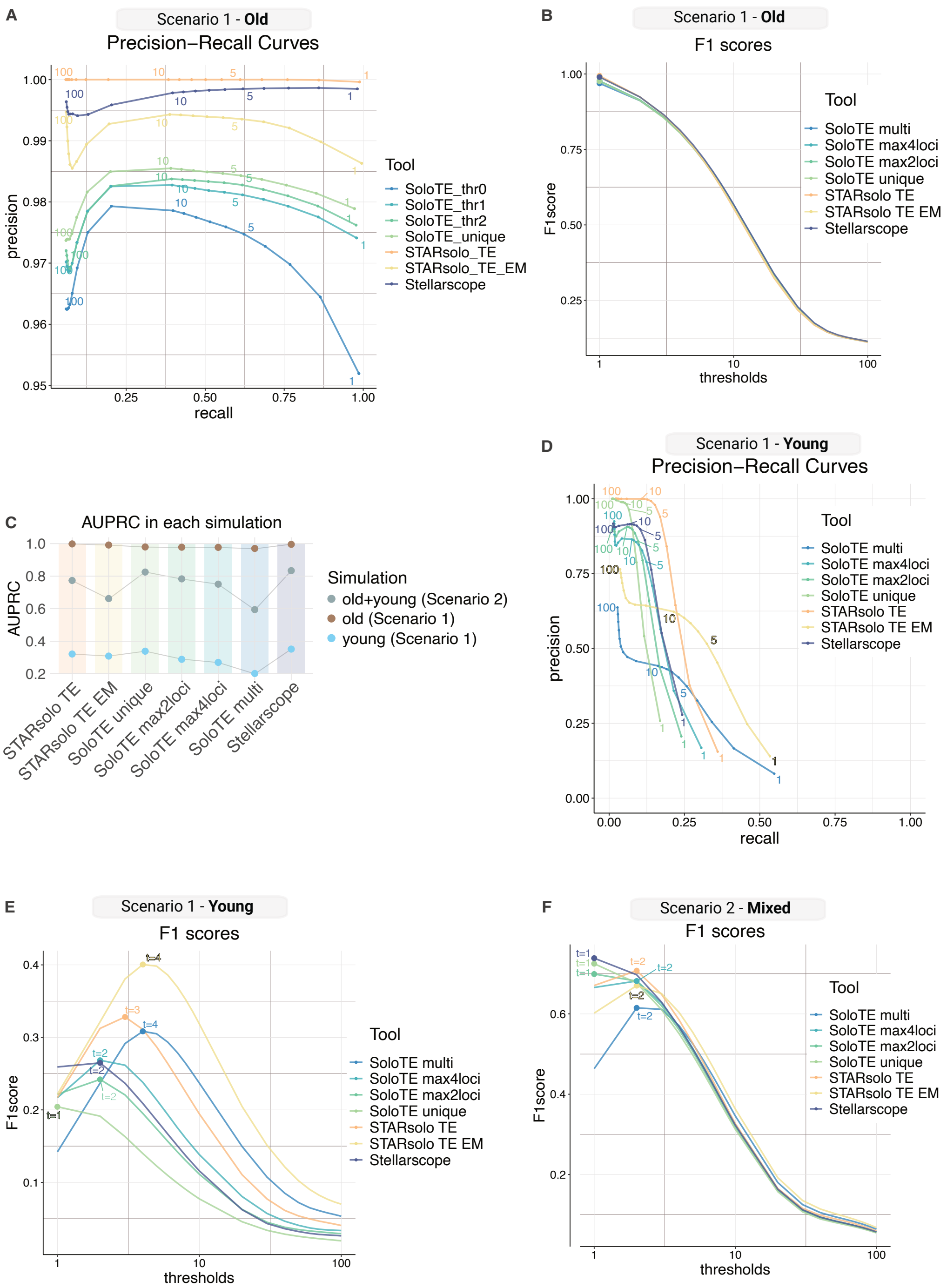


Figure S4: **A** PR curves obtained by varying count threshold for locus detection (1–10 with step 1 and 20–100 with step 10) in the Scenario 1 old simulation. Curves are computed separately for each TE quantification method. **B** F1 score of TE detection obtained after varying count threshold for locus detection (1–10 with step 1 and 20–100 with step 10) in the Scenario 1 old TE simulation. The labeled dots indicate the count threshold which resulted in the highest F1 score for each tool. The x-axis is log10-scaled. **C** AUPCR obtained by varying the count threshold from 1 to 100 for locus detection with each tool, in each of the simulations from Scenario 1 and 2. **D** PR curves obtained by varying count threshold for locus detection (1–10 with step 1 and 20–100 with step 10) in the Scenario 1 young simulation. Curves are computed separately for each TE quantification method. **E, F** F1 score of TE detection obtained after varying count threshold for locus detection (1–10 with step 1 and 20–100 with step 10) in the Scenario 1 young (E), and the mixed Scenario 2 (F) simulations. The labeled dots indicate the count threshold that resulted in the highest F1 score for each tool. The x-axis is log10-scaled.

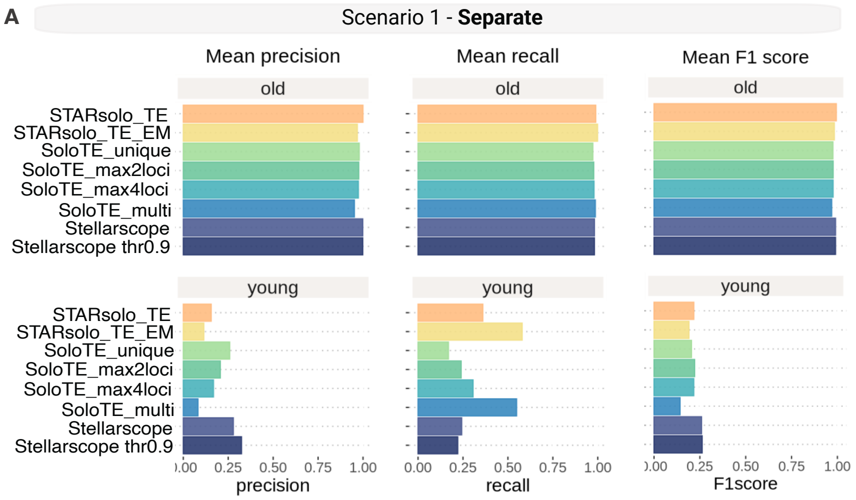


Figure S5: **A** Precision, recall and F1 score computed per cell and then averaged across cells, using binarized TE count matrices. Results are shown for simulations with old loci (top row) and with young loci (bottom row), including the results of Stellarscope after applying a threshold of 0.9 on posterior probability scores.

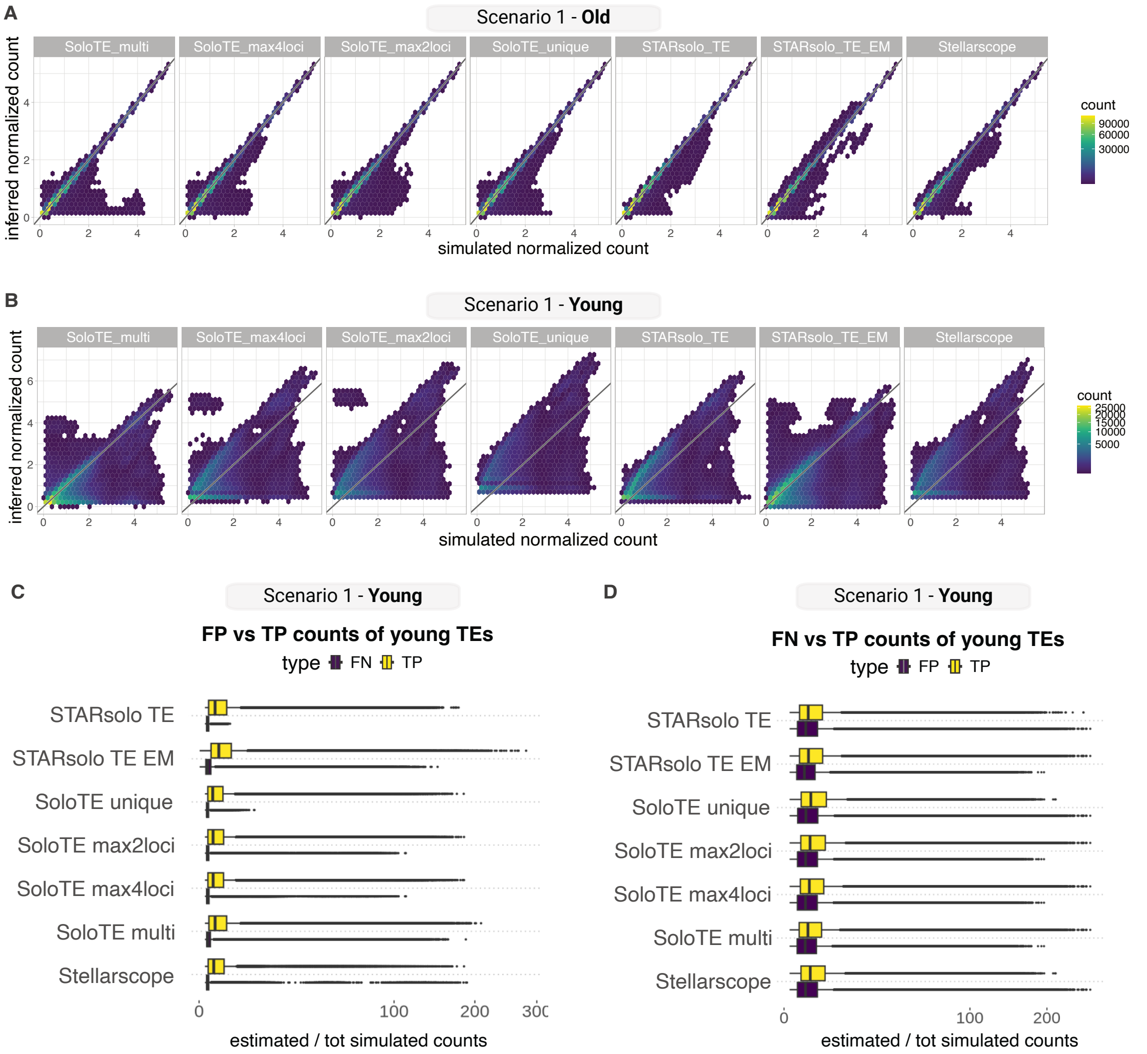


Figure S6: **A**,**B** Density plots comparing simulated (x-axis) and estimated expression levels of old (A) and young (B) TEs. Log-normalized counts. **C** Boxplots showing the distribution of expression levels of TPs and FPs. Raw counts were normalized by the total number of simulated counts per cell. **D** Boxplots showing the distribution of expression levels of TPs and FNs. Raw counts were normalized by the total number of simulated counts per cell.

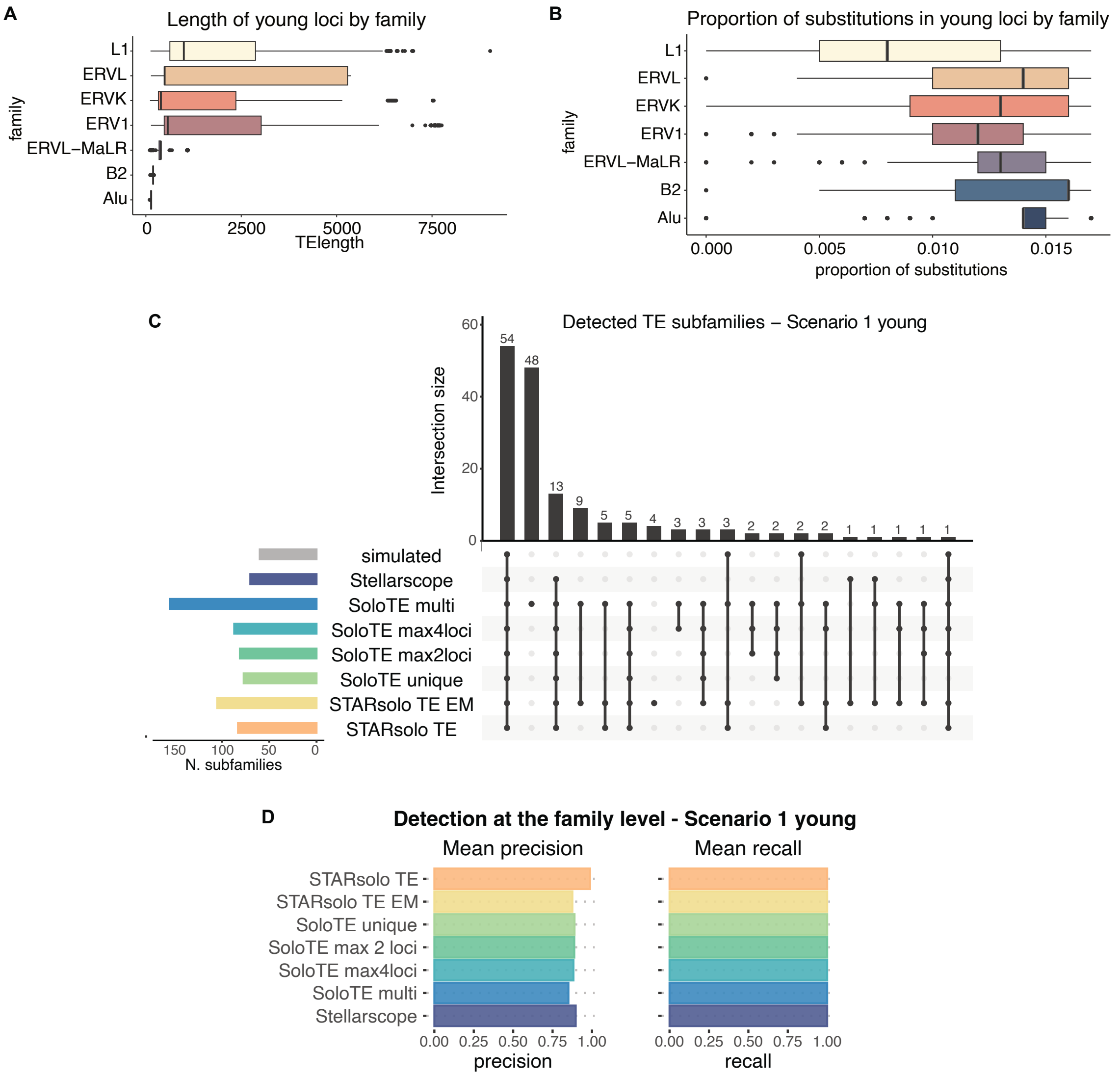


Figure S7: **A** Boxplot of length in base pairs of young loci included in the simulations, stratified by family. **B** Boxplot of the proportion of substitutions reported in the RepeatMasker annotation for young loci included in the simulations, stratified by family. **C** Upset plot of sets of TE subfamilies present in the old TE simulated matrix (grey bar) and detected as expressed by each tool. **D** Precision and recall computed per cell and then averaged across cells, using subfamily-aggregated, binarized TE count matrices. Results are shown for simulations with young loci.

Table S2: Information on publicly available datasets used in our benchmarking.

| **Cell type** | **Species** | **Technology** | **Number of cells passing QC** | **Number of input reads** |
|---|---|---|---|---|
| ESCs | Mouse | 10X Chromium Single Cell 3' v3 | 2,194 | 215,105,630 |
| PBMCs | Human | 10X Chromium Single Cell 3' v3 | 8,243 | 784,064,148 |
| Olfactory Mucosa | Mouse | 10X Chromium Single Cell 3' v3 | 34,871 | 541,486,729 |

Table S3: Marker genes used for mESC cell type annotation.

| **Cell state** | **Marker genes** |
|---|---|
| Pluripotent | Zfp42, Sox2, Nanog |
| 2CLC | Zscan4a, Zscan4c, Zscan4d, Zscan4e |
| Differentiated | Gata6, Sox17, Sox7 |

Table S4: Marker genes used for PBMC cell type annotation.

| **Cell type** | **Marker genes** |
|---|---|
| B cells | CD79A, MS4A1 |
| Dendritic cells | FCER1A, CST3, LYZ, LGALS3 |
| $CD14^+$ Monocytes | CD14, LYZ, LGALS3, S100A8 |
| $FCGR3A^+$ Monocytes | FCGR3A, LYZ, CST3 |
| Natural Killer cells | GNLY, NKG7 |
| $CD4^+$ Memory T cells | IL7R, S100A4 |
| $CD4^+$ Naive T cells | IL7R, CCR7 |
| $CD8^+$ T cells | IL7R, CD8A, CD8B |
| Platelets | PPBP |

Table S5: Marker genes used for mouse olfactory mucosa cell type annotation.

| **Cell type** | **Marker genes** |
|---|---|
| Globose basal cells (GBC) | Ascl1, Kit |
| Immediate neuronal precursors (INP) | Neurod1, Gng8 |
| Immature olfactory sensory neurons (iOSN) | Gng8, Gap43 |
| Mature olfactory sensory neurons (OSN) | Omp, Adcy3 |
| Horizontal basal cells (HBC) | Trp63, Krt5 |
| Microvillar cells (Mv1) | Trpm5 |
| Microvillar cells (Mv2) | Ascl3, Cftr |
| Sustentacular cells (Sus) | Cyp2g1 |
| Immune cells | Gypa |
| Blood cells | Ptprc |
